# Perceptual tri-stability, measured and fitted as emergent from a model for bistable alternations

**DOI:** 10.1101/2024.05.24.595419

**Authors:** Jiaqiu Vince Sun, Zeyu Jing, James Rankin, John Rinzel

## Abstract

The human auditory system in attempting to decipher ambiguous sounds appears to resort to perceptual exploration as evidenced by multi-stable perceptual alternations. This phenomenon has been widely investigated via the auditory streaming paradigm, employing ABA_ triplet sequences with much research focused on perceptual bi-stability with the alternate percepts as either a single integrated stream or as two simultaneous distinct streams. We extend this inquiry with experiments and modeling to include tri-stable perception. Here, the segregated percepts may involve a foreground/background distinction. We collected empirical data from participants engaged in a tri-stable auditory task, utilizing this dataset to refine a neural mechanistic model that had successfully reproduced multiple features of auditory bi-stability. Remarkably, the model successfully emulated basic statistical characteristics of tri-stability without substantial modification. This model also allows us to demonstrate a parsimonious approach to account for individual variability by adjusting the parameter of either the noise level or the neural adaptation strength.

## Introduction

Imagine yourself resting in a park at dusk, eyes closed, and enveloped by different sounds. Remarkably, your brain instinctively organizes these sounds into distinct sources, allowing your mind to drift effortlessly from one source to another. The auditory system has an innate capability to discern and track multiple sound streams, allowing multiple perceptual interpretations in cases of ambiguity. This has been conceptually framed as auditory scene analysis (Bregman, 1990), and experimentally investigated using sequences of pure tones consisting of repeating ABA_ triplets (Noorden, 1975). Here, “_” represents a silent gap. Such a tone sequence aligns with multiple perceptual interpretations (percepts), and only one of them dominates at a time. During minutes-long presentations, the dominant percept switches stochastically after the initial transient build-up stage. Most studies considered only two percepts (Schwartz et al., 2012; Szabó et al., 2016): an “integrated” stream (ABA_ABA_…) and two “segregated” streams (A_A_A_A_… and B B) (Figure 1A, B, green and purple).

**Figure 1.**
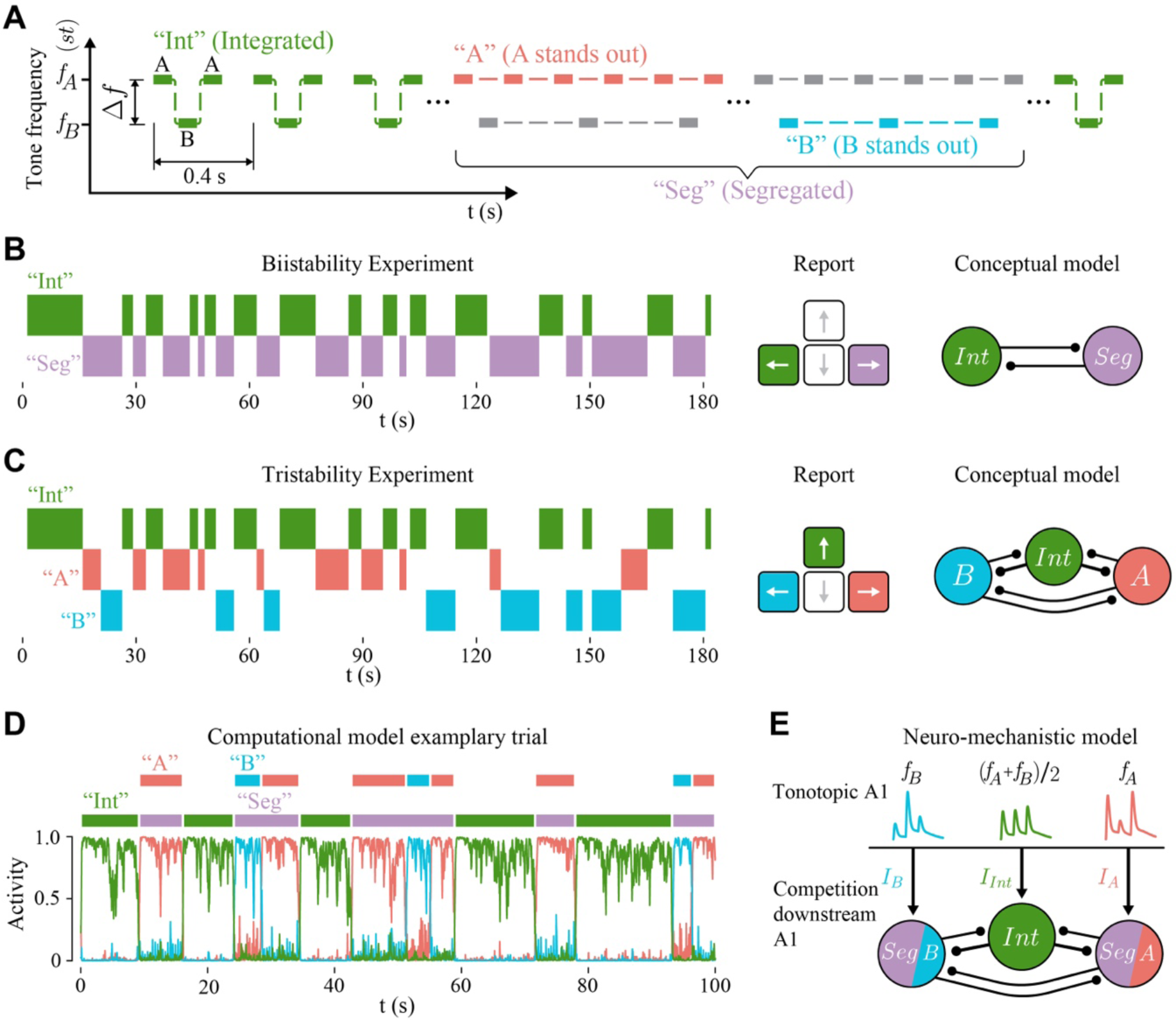
Multi-stability paradigms. **A**, In a single trial (3 mins), a participant listens to two interleaved sequences, one of A tones and the other of B tones; the organization appears as tone triplets ABA_ABA_ The repetition rate of A tones is twice that of B tones. In the tri-stable paradigm, a participant reports continuously on the current perception as one of three interpretations: as one integrated stream ABA_ABA_… (green), or as two segregated streams with either an A_A_A_A_… sequence standing out (A percept, red) or a B B sequence stands out (B percept, blue). **B,** In the bi-stable paradigm, only two stochastically varying percepts (left) are considered (“Int” and “Seg”). A participant reports the current percept by holding down one or the other of two keys (middle). A typical conceptual model of bi-stability involves two percept-encoding units that compete for dominance (right). **C,** The current study features perceptual tri-stability illustrated in **A**. A participant reports (left) three stochastically varying percepts (“Int”, “A”, and “B”) by holding down one of three keys at a time (middle). A typical model of tri-stability involves three units that compete for dominance (right). D, the simulated rate time courses of the three populations from the neuro-mechanistic model shown in **E.** Only one population is dominant at a time. The on and off of the Int unit correspond to “Int” and “Seg” percepts when the model is applied to perceptual bi-stability. **E,** Our tone-based, neuro-mechanistic model has two stages, the sensory stage and the perceptual-competition stage. The tone-evoked activities in the tonotopically-organized sensory stage (primary auditory cortex A1) provide input to the competition stage in which three groups of E and I neuronal populations compete via mutual inhibition to determine perceptual dominance. This 2-stage, 3-unit model was developed and applied to modeling bi-stable perception: “Int” (green) and “Seg” (purple).

However, perceptual interpretations beyond this dichotomy were mostly neglected, despite one study (Denham et al., 2014) that identified as many as six percepts in the same ABA_ streaming paradigm, and a corresponding conceptual model (Winkler et al., 2012). Admitting the difficulty in training participants to reliably identify six percepts, we believe tri-stability is a good starting point to constrain models of perceptual multi-stability that admits more than two percepts. An intuitive extension is to further distinguish foreground versus background stream within the “segregated” (Figure 1A, red and blue) category, in addition to the “integrated” percept.

Current models of auditory perceptual multi-stability (Figure 1B, right), to the best of our knowledge, have yet to be constrained by situations that allow for more than two interpretations. In contrast, tri-stability has been explored in visual perception. For instance, Huguet et al. (2014) developed and tested models of tri-stability of viewing dynamic plaids, overlapping drifting gratings. The experimental task and the model involve distinguishing foreground and background when separated streams are being perceived. However, in audition, this foreground-background switching tends to be more variable and difficult to restrict even after introducing a strong biasing cue (Hupé & Pressnitzer, 2012). Additionally, the sensory responses for the triplet tones involve onset/offset transients, contrasting with the typically maintained sensory responses to visual plaids. The ongoing transients pose a challenge for a biologically-constrained model of auditory perception that should sustain a percept over several triplets (Rankin et al., 2015). Furthermore, the differing repetition rates of the two tones in auditory perception introduce an asymmetry in the segregated percepts, an aspect not observed in visual plaids.

This study endeavors to investigate auditory tri-stability through our sensory-based, physiologically plausible neuro-mechanistic model (Rankin et al., 2015). The model has been applied to accurately reproduce main features of auditory bi-stability dynamics. Intriguingly, the model incorporates tri-stability features, although its original interpretations focused on bi-stability. The model encompasses two hierarchical and tonotopically organized stages: one for sensory processing (like primary auditory cortex, A1) and another for neural competition, downstream of A1. The competition stage involves three neuronal subgroups selective primarily to tones A or B or to both; dominant activity by the central unit is categorized as the representation of “integrated”, while dominance by either peripheral unit is classified as “segregated”. In the current study, we find in segregation that one or other of the two peripheral units represent A and B tone streams is dominant, representing as foreground (“A” or “B”) percepts, respectively.

Our model adopts a dynamic systems approach to model biased competition (Laing & Chow, 2002; Wilson, 2003; Moreno-Bote et al., 2007; Shpiro et al., 2009; Mill et al., 2013; Meso et al., 2016; Little et al., 2020). It stands apart from other models, such as those based on evidence accumulation (Barniv & Nelken, 2015; Cao et al., 2016; Nguyen et al., 2020) and Bayesian inference (Gershman et al., 2012), by its sensory-rooted design and foundations in neurophysiological research on auditory streaming (Fishman et al., 2001; Gutschalk et al., 2005; Micheyl et al., 2005).

In this study, we gathered data from people as they experienced tri-stable perceptions during extended presentations of the ABA_ tone triplet. We varied the frequency separation of the two tones, measured in semitones (st) and denoted as Δf, across trials. Large Δfs of ±13 st were selected to foster segregated percepts and enhance tri-stability. Our findings indicate that the model adeptly replicated the statistical patterns of auditory tri-stability at both individual and group levels, achieving this without substantial modifications. The notable individual differences observed in our study are attributable to variations in neural noise or adaptation among participants.

The paper unfolds as follows: The **Material and Methods** section explains the stimuli and experimental procedures. In the Theory and Calculation section, we first revisit our computational model for bi-stability, then describe numerical simulations and model fitting. In the **Results** section, we start by presenting new results and insights from our tri-stability psychophysics experiments. Next, we detail how the model was recalibrated to explicitly accommodate tri-stability, and how the model simulations faithfully reflect both individual and collective behavioral patterns. The **Discussion** section explores some serendipitous experimental observations and discusses the role of noise, adaptation, and attention in our model. It also covers different types of models of perceptual multi-stability and situates our tri-stability findings in a broader landscape. Notations: “A”, “B” and “Int” in quotation marks represent distinct percepts.

## Material and methods

### 1. Stimuli

The stimuli consist of interleaved sequences of A tones and B tones, whose organization appears as tone triplets ABA_ABA_… (**Figure 1A**) The repetition rate of A tones is twice that of B tones. Each tone is 100-ms long including 5-ms Cosine squared rise and fall. Each triplet is separated by a silence of the same length indicated by ‘_’; therefore, each triplet is 400 ms long. The A tone and B tone are apart in pitch by Δ*f* semitones (st). A positive Δ*f* indicates that the A tone is higher than the B tone, while a negative Δ*f* indicates the opposite. During a 3-minute-long trial, the tone sequence is played binaurally through Etymotic earphones at 65 dB SPL.

To examine the model’s explanatory power in a wider scope, we selected four Δ*f*s: +2, +5, and ±13 st as our experimental conditions. The frequency difference of the two tones (Δ*f*) and the presentation rate (PR) are the major parameters that influence the dominance statistics and switching dynamics (Noorden, 1975). In our experiment, Δ*f*=+2 served as part of the data sanity check as it should be dominated by integrated percept. Δ*f* = +5 was the equidominance point for bi-stability (Int and Seg) in previous studies where the same model in this paper was applied (Rankin et al., 2015, 2017; Byrne et al., 2019; Rankin & Rinzel, 2022). Large Δ*f*s (13) would promote the two segregated percepts in tri-stability. Additionally, we introduced a novel experimental condition by inverting the frequencies of A tone and B tone to low-high-low (Δ*f* = −13), thereby broadening the test conditions, and challenging the model further. The orders of the high and low frequencies and the saliency of the tones have been mostly assumed to be irrelevant in the literature. However, (Denham et al., 2014) found that the frequency order affects the dominance statistics and speculated about an influence from tone saliency. Therefore, while the standard approach is to roam the tone’s absolute frequencies in an arbitrary range, we carefully arranged the frequencies according to the equal loudness contour to constrain tone saliency while minimizing potential adaptation. Finally, we fixed PR at 10 Hz because a fast PR would also promote segregated percepts.

### 1. 2. Experimental procedure

17 subjects (14 females) with a mean age of 25 years participated in the experiment and were reimbursed for their time at a $10 hourly rate. The procedures were approved by the University Committee on Activities Involving Human Subjects at New York University (IRB_FY2016-310). All participants provided written informed consent.

Anticipating increased complexity and variability associated with tri-stability versus bi-stability (Hupé & Pressnitzer, 2012), we arranged two sessions in the experiment to collect enough data to look at individual subjects. The first session has four conditions (Δ*f* = +2, +5, ±13 st) with four trials for each condition while the second session has three conditions (Δ*f* = +5, ±13 st) with four trials for each condition. Each pair of frequencies appear no more than twice in a session. After completion of the first session, four of the participants did not pass the sanity check because they in general made very few perceptual switches in different conditions. 13 participants were told to do the second session on different days, while 11 of them came back. All participants had normal hearing verified by a hearing screening. The participant sat comfortably in a soundproof cabin.

At the beginning of the first sessions, participants first went through a hearing screening. Their sound detection thresholds at a range of frequencies were measured. Then, they went through a brief training. It started at extreme Δ*f*s to familiarize participants with “Int” with Δ*f* = +2 and “A” and “B” with Δ*f* = +13. Next, they went through several 1-minute mock-up trials with different Δ*fs*. They were instructed to report the current perception by holding down the corresponding key–Up for integration, Left for the low-pitch sequence standing out, and Right for the high-pitch sequence standing out (Figure 1C, middle). The data generated in a 3-minute long trial would look like Figure 1C, left. Participants were told not to intentionally control the perception but to let it vary spontaneously. The training ended after they felt confident in reporting the perception and switching the key immediately when the perception had changed.

After the first session, we performed data sanity checks by applying a few criteria to the participants. First, we ensured that the proportion of time of Int was greater than the other two percepts respectively at Δ*f* = +2. Second, we ensured that subjects at least reported on average two perceptual switches in a trial. Four subjects who did not meet the second criterion were informed not to join the second session. Another two subjects missed the second session for personal reasons. In total, 11 subjects participated in the second session, whose data was included in the analysis and model fitting.

## Theory and calculation

### 1. Basic model structure

Our model describes the dynamic perceptual alternations by adopting a firing rate-based dynamical systems framework. The dominance of a percept at a given time is instantiated by a winner-take-all competition among three percept-encoding populations of excitatory and inhibitory (E and I) neurons (Figure 1C, right), facilitated by mutual inhibition, slow adaptation, noise–features that have been widely used to explain perceptual bi-stability (e.g, Laing & Chow; Wilson; Shpiro et al., 2007, 2009; Huguet et al., 2014). Inclusion of moderately slow local recurrent excitation enables percept persistence across several triplets (Rankin et al., 2017). The model has been adapted to explain the perceptual build-up stage (Rankin et al., 2017), stimulus perturbations (Rankin et al., 2017), dynamic entrainment (Byrne et al., 2019), and attentional gain control (Rankin & Rinzel, 2022).

The model assumes two stages of processing (Figure 1E; See Figure 2A for more details). The pre-competition stage employs dynamic inputs directly connected to sensory features as represented by neuronal responses from the primary auditory cortex (A1). The inputs’ spatiotemporal profiles are based on A1 responses to interleaved A and B tones in primate electrophysiology studies (Fishman et al., 2001, 2004). The competition stage features a tonotopic organization with three populations/units of E-I neurons, assumed to be downstream A1. The two peripheral populations (A unit and B unit) receive input from A1 regions where *f_A_* or *f_B_* is preferred. A third population (AB/Int unit) receives input from tonotopically intermediate A1 locations, roughly centered at (*f_A_* + *f_B_*) / 2. In the competition stage, each population/unit (Figure 2B) has instantaneous self- and mutual inhibition, receives inputs from A1, receives additive noise from A1 and other non-local sources, has recurrent NMDA excitation, and undergoes slow adaptation (timescale 1.4 s). The inhibition strength from Int unit is twice the inhibition strength from A unit and B unit. The model details related to tri-stability are further described in Results 2. The extra mechanistic details can be found in Rankin et al. (2015), with a few corrections in Byrne et al. (2019). See Model Equations at the end of the Supplemental Information and Supplementary Table for parameter values.

**Figure 2.**
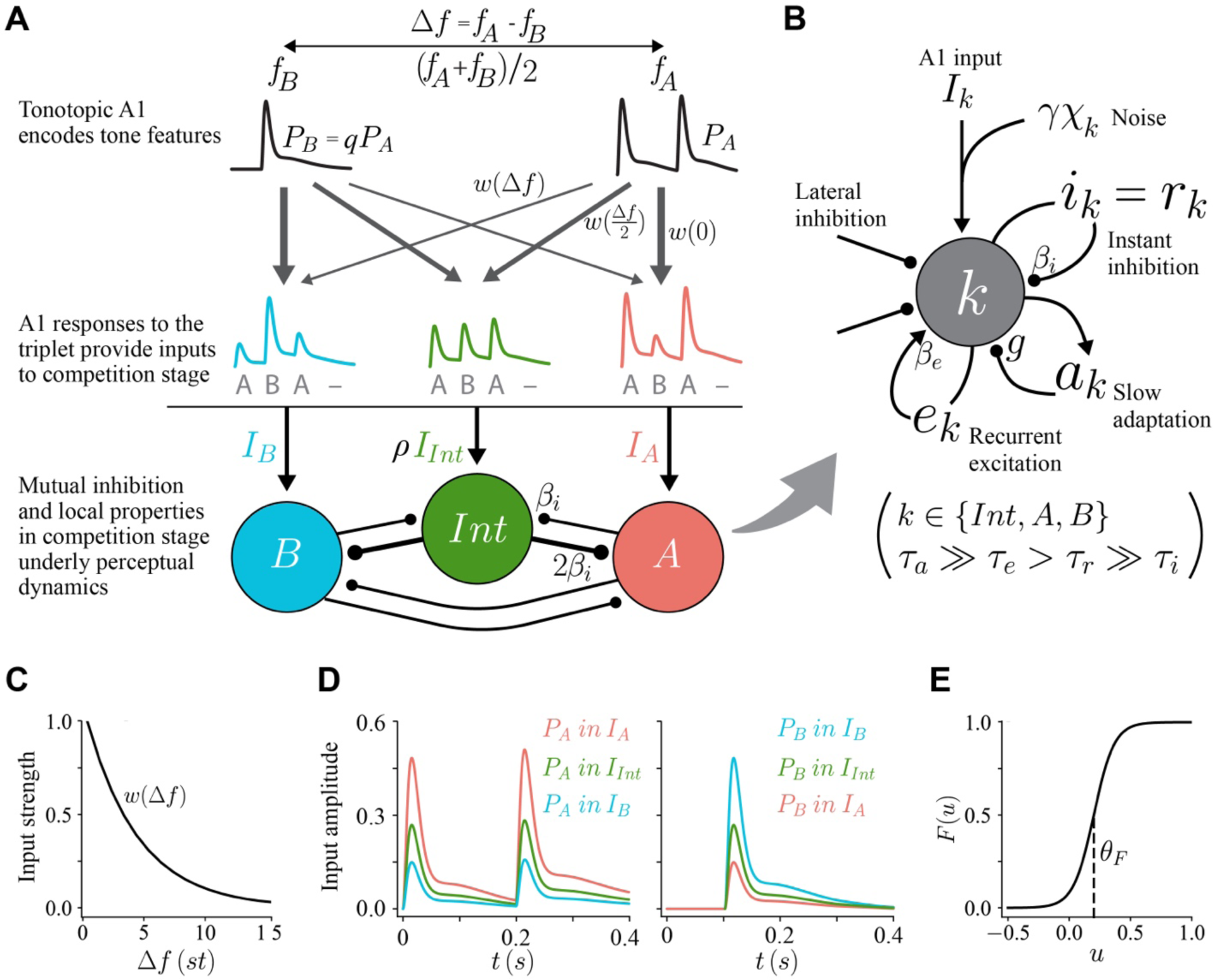
Two-stage neuro-mechanistic model. **A**, Model schematic with two stages (Rankin et al., 2015) – tonotopic primary auditory cortex (A1) that only encodes stimulus features and percept-encoding populations downstream A1 that compete for dominance. We include a parameter q to reflect stimulus saliency (e.g., if a higher pitch increases saliency). q is the ratio of the peak amplitudes of the response PB to B tone and P_A_ to A tone at their frequency-preferred locations respectively in A1. Tones generate across A1 onset-then-plateau responses, whose amplitudes (as inputs to the competition stage) scale with the input function *w*(Δ*f*) shown in **C**. For a triplet of ABA_ tones, the pitch difference Δ*f* in semitones (st) dictates the input profiles of *I*_int_, *I*_A_, and *I*_B_ (also shown in **E**), with *I*_int_ mostly contributed by areas centered at (*f*_A_ + *f*_B_)/2. **B.** The percept-encoding populations (Int, A, and B) have instantaneous self- and mutual inhibition, receive inputs from A1 and additive noise from A1 or other non-local sources, have recurrent NMDA excitation, and undergo slow adaptation (timescale 1.4 s). **E**, The sigmoid activation function for each population: steady response for steady input.

Criteria based on population firing patterns classify perceptual interpretations (Figure 1D). Despite that the three-population model naturally allows for tri-stability, it was applied previously only to perceptual bi-stability. That is, (Figure 1B, left and middle) participants were instructed to report on perceiving the “integrated” (“Int”) or the segregated (“Seg”) without distinguishing between stream A being in the foreground (“A”) and stream B being in the foreground (“B”). In the model’s original application (Figure 1D, E), the current perception is “Int” when Int unit is on and “Seg” when Int unit is off.

### 2. Numerical simulation and readout of percepts

We conducted simulations using Python, employing a standard Euler-Murayama time-stepping method with a step size of 2 ms, which corresponds to one-fifth of the fastest timescale parameter in our equations (τ_r_ = 10 ms). Using a smaller time step did not noticeably alter the results.

We extracted dominant percepts from the population firing rate time courses, namely *r*_B_, *r*_int_, and *r*_A_. We first smoothed the firing rates with a 50-ms-wide moving average filter. Then, we categorized the dominant percept based on the highest firing rate at each point in time, and combined neighboring time points with the same percepts. The behavioral task would require a minimum duration for identifying a perceptual switch. As such, we set a 500-ms threshold, and we merged fragmented percepts (lasting less than the threshold) with their neighboring percepts. We processed a sequence of contiguous fragmented percepts as a single cluster. The direction of merging followed an intuitive rule: if the most dominant percept in the cluster is the same as the previous percept, the cluster will be merged backward into the previous percept. Otherwise, it will be merged forward into the next percept. If no percept in the cluster exceeds 60%, the cluster will be labeled as ambiguous (“Amb”).

### 3. Fitting the model for individual subjects by tuning the noise or the adaptation parameter

We fitted two sets of parameters–noise, γ and adaptation, *g*–to individual subjects’ mean durations of different percepts in various conditions. The two parameters were selected based on explorative simulations (Results are shown in Supplementary Figure 1). We only used the mean duration to calculate the fitting error for several reasons. First, we observed substantial individual differences in the mean durations of the experimental results. Second, the mean durations exhibited monotonic dependency on the parameters of interest for a given Δ*f* in our exploratory simulations (Supplementary Figure 1). Moreover, fitting for the mean durations allowed for flexibility in selecting percepts and Δ*f*s when computing the loss. While the proportions of time of different percepts at a given Δ*f* are interrelated, their mean durations are not. As such, conditions in which some participants did not yield enough percepts (30 occurrences) for a reliable estimation were excluded from the fitting (Supplementary Figure 2). The percepts used in the fitting contained “Int” and “A” at Δ*f* = +5, “A” at Δ*f* = +13, and “A” and “B” at Δ*f* = −13.

We fit the parameters (γ or *g*) for the 13 eligible participants and for different absolute values of Δ*f*s (5 and 13) respectively. We ran a single grid search for each parameter on a high-performance computing system (HPC). For each parameter value, we ran 12 trials of six-minute simulations, totaling 4320 s. We excluded the first (buildup) and last (incomplete) percepts from these trials and calculated the statistics for all perceptual categories and Δ*f*s. In cases where simulations resulted in exceptionally long percept durations (due to low noise or weak adaptation), we carried out additional trials of simulations to ensure that the least frequent perceptual category occurred at least 100 times.

At the group level, we fine-tuned the tone-response gain (*q*) and the attentional gain (ρ). We maintained a fixed value of 0.94 for *q* across simulations, while ρ was 0.9 when fitting noise and 1.0 when fitting adaptation. Intriguingly, fitting adaptation was insensitive to the value of ρ. Therefore, we keep ρ = 1.0 as a parsimonious account. the Int unit was not more or less biased than the A and B units.

## Results

We organize the results as follows. **Part 1, Behavior experiments,** presents the behavioral results of our tri-stability experiment. **Part 2, Modeling tri-stability,** explains the modification we introduced to account for tri-stability. **Part 3, Model simulations reproducing the individuals’ and the group’s key statistical characteristics,** compares simulations with experimental results at different levels.

### 1. Behavior experiments

We examined both the group-average trends and the individual differences in three statistic indicators–the proportion of time, mean durations, and probability of occurring of three percepts (“Int”, “A”, “B”) under three conditions (Δ*f* = +5, ±13). The proportion of time is the cumulative duration of a percept in a trial relative to the trial’s total length. The mean duration reflects the average time a percept persists, while the probability of occurring is defined as the frequency of a specific percept’s appearance relative to the total number of percept occurrences. For clarity, we only present the proportion of time and mean duration results in this part (detailed findings regarding the probability of occurring are presented in Results subsection 4. Model simulations…).

At the group level, the behavioral patterns observed in tri-stability were broadly consistent with bi-stability, albeit with some distinct differences. As Δ*f* increased from +5 to +13, we observed a decline in the proportion of ‘Int’ from 44% to 17% (*t*(10) = 7.55, *p* < 0.0001, *d_z_* = 2.72). Concurrently, the proportion of “A” rose from 43% to 63% (*t*(10) = −8.70, *p* < 0.0001, *d_z_* = 1.82), and the proportion of “B” from 13% to 20% (*t*(10) = −2.72, *p* = 0.043, *d_z_* = 0.81) (Figure 3A, left panel). Unlike the proportion of time, the mean duration of “Int” decreased only marginally with an increased Δ*f* (*t*(10) = 1.85, *p* = 0.187, *d_z_* = 0.5) (Figure 3A, right). These trends were in line with previous findings in auditory bi-stability (Billig et al., 2018; Rankin et al., 2015; Rankin & Rinzel, 2022). Notably, at Δ*f* = +5, an equidominance between “Int” and “A” emerged–a departure from the bi-stable equidominance previously reported between “Int” and “Seg”. If we consider consecutive “A” and “B” as a combined “Seg”, this pseudo “Seg” percept occupied a greater portion of time than “Int” (*t*(10) = −3.01, *p* = 0.026, *d_z_* = 1.82, Figure 3B, right). We attributed these findings to task effect: the tri-stability task likely intensified the focus on distinguishing segregated percepts (as detailed in the Modeling tri-stability section). Furthermore, the less definitive and more challenging foreground-background dissociation in the tri-stability task may have boosted the attention given to segregated percepts due to the heightened task difficulty.

**Figure 3.**
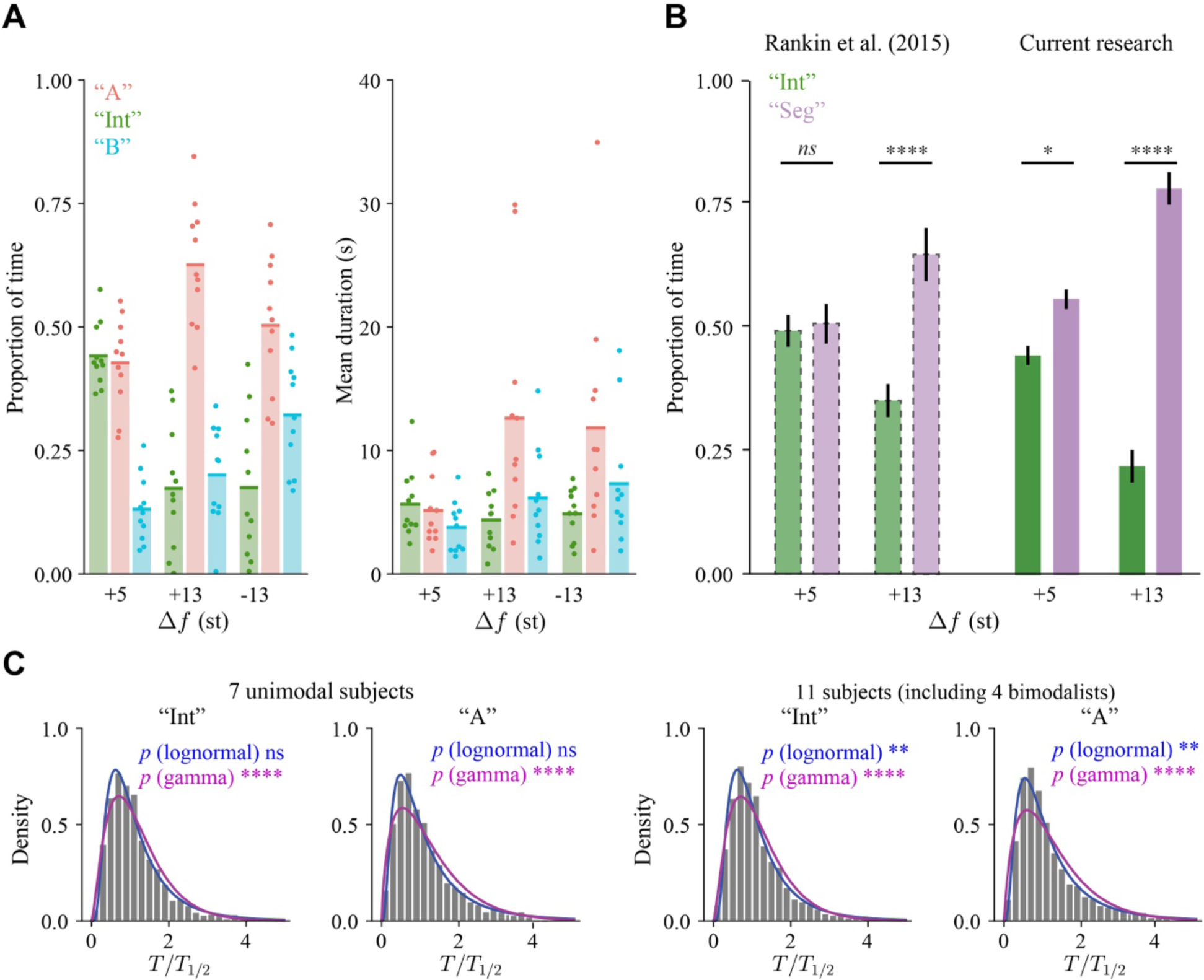
Results of the behavioral experiment. **A**, Group average (shadings and lines), and individual subjects’ (dots) proportion of time (left) and mean durations (right) of the three percepts (color coded). _For_ the group average, the proportion of time and mean duration show similar patterns for given Δ*f* but divergent patterns across Δ*f*. The proportion of time for “Int” drops substantially from Δ*f* = +5 to Δ*f* = ±13. Interestingly, individuals’ mean durations at Δ*f* = ±13 are exceptionally variable, which suggests fitting a model to each individual’s mean durations. **B**, The task-induced change of bi-stable equiponderance in the current task (right panel) compared with Rankin et al. (2015) (left panel). In the current study, pseudo “Seg” percepts were created by merging consecutive “A” and “B” percepts. The proportion of time in Rankin et al. (2015) was created by interpolating from a published figure. The ratios of the proportion of “Seg” to “Int” are higher in the current study than previously reported for all the Δ*f*s tested. Δ*f* = +5 is no longer an equidominance point between “Int” and “Seg” in the current study. **C**, Histograms of median-normalized durations aggregated over Δ*f*s and subjects are well approximated by lognormal distributions for a subpopulation of seven whose have unimodal histograms (upper row). Histograms of eleven subjects including four bimodal ones are no longer lognormal. None of the histograms are approximated by a gamma distribution. All eleven subjects are included in behavioral analysis and model fitting. * indicates *p* < 0.05, ** *p* < 0.01, *** *p* < 0.001, and **** *p* < 0.0001, ns nonsignificant (which means the histogram is not significantly different from the distribution in the KS test).

Switching the two tones’ frequencies (from high-low-high to low-high-low) caused readjustment of “A” and “B”, while “Int” remained largely unaffected (Figure 3A). When Δ*f* shifted from +13 to −13, the proportion of “A” decreased from 63% to 50% (*t*(10) = 2.65, *p* = 0.024, *d_z_* = 0.94), and the proportion of B increased from 20% to 32% (*t*(10) = −3.41, *p* = 0.013, *d_z_* = 1.14), with no significant change in the proportion of “Int” (*t*(10) = −0.07, *p* = 1.90, *d_z_* = 0.01) (Figure 1A, left). These observations are consistent with our choices of tone frequencies, where the higher frequency tone was more prominent than the lower one (see Methods section). However, the effect of tone frequency reversal in our experiment was subtler compared to a previous study on auditory multi-stability that accommodated more than three interpretations (Denham et al., 2014). Three one-way repeated measure ANOVAs revealed that Δ*f* significantly influenced the proportion of time for all percepts (see Table 1 in Appendix A). While the overall trend for mean durations paralleled that of proportion of time, the effect of Δ*f* on “Int” was not significant (*F*(2,20) = 1.32, *p* = 0.291, η^’^=0.051). Moreover, the mean duration of “A” was considerably longer at Δ*f* = +13 (12.6 s) than at Δ*f* = +5 (5.1 s) (*t*(10) = −3.47, *p* = 0.012, *d_z_* = 1.09). In summary, switching tones’ frequencies affected the proportion of time more noticeably than the mean duration.

Beyond group-level findings, we noted substantial individual variability, particularly regarding the most dominant percept, “A”, at larger Δ*f*s. The mean duration of “A” varied widely, from 2.5 s to 29.9 s at Δ*f* = +13, and from 1.9 s to 34.9 s at Δ*f* = −13 (Figure 3A, right). This variability is underscored by the coefficient of variation (CV) of 0.70 or higher. These large discrepancies among subjects underscore the necessity of tailoring our model to each individual participant.

To get an overview of the distributions of percept durations, we aggregated the normalized durations over the 11 subjects and three Δ*f*s (+5, ±13) for each percept (Figure 3C, right). Interestingly, the resulting histograms of these normalized durations did not conform to either gamma or log-normal distributions. Further analysis revealed that four subjects exhibited at least one bimodal distribution among the three percepts (see Supplementary Figure 3). When aggregated only from the seven subjects with unimodal distributions, the histograms aligned with log-normal distributions (Figure 3C, left). The histograms of aggregated data for each combination of Δ*f* and percept confirmed the results (Supplementary Figure 4 and 5). However, subjects with bi-modal distributions of percept durations were rarely reported in previous research. Despite this, we included these four bimodal subjects in our main analysis as our model-fitting approach embraces, rather than overlooks, individual differences. While we aimed to fit the model to individual subjects’ summary statistics, the underlying causes of bimodality were not a focus of this study.

### 2. Modeling tri-stability

The current study examines and builds upon the model proposed by Rankin et al. (2015), with the specific goal of accounting for auditory tri-stability. The dominance of either peripheral unit was classified as segregated (Figure 1D) in previous applications of this model (See methods for the classification criteria). The current study assumes that the units in the competition stage correspond respectively to the three percepts–“B”, “Int”, and “A” (Figure 1E). In addition to this reinterpretation, we augmented our model with two new parameters, reflecting tone saliency and task-induced attentional gain respectively. The two parameters readjust A1 response profiles to the triplets and the inputs to the competition stage. Our goal was to find minimal yet effective modifications that would enable the model to adeptly fit individual participant’s data.

The tone-response gain parameter *q* (Figure 2A, top row) determines the ratio of the amplitudes (*P*_A_, *P*_B_) of A1 responses to the higher-pitch tone and the lower-pitch tone, dictated by the tones’ saliency. *q* = *P*_A_ / *P*_B_ when Δ*f* > 0; *q* = *P*_B_/ *P*_A_ when Δ*f* < 0. In the pre-competition stage (A1), the two tones evoke across tonotopic locations periodic onset plateau responses (Micheyl et al., 2005) whose amplitudes decay exponentially (*w*(Δ*f*)) as moving away from their frequency-preferred locations (Figure 2C, D). Additionally, the parameter *q* influences the undulation degree in the A1 output profiles in response to the ABA_ triplet, as illustrated in the middle row of Figure 2A. The experimental stimuli (Supplementary Figure 6) were designed within the constraint of 0< *q* <1 (see Methods).

The attentional gain parameter ρ scales the total input to the Int unit (*I*_int_) in the competition stage (Figure 2A bottom), relative to the inputs to A unit and B unit (*I*_A_ and *I*_B_). It reflects the attentional bias towards the two segregated percepts: 0 < ρ < 1. The tri-stability task itself requires dissociating “A” and “B”, which could be more difficult than distinguishing “Int” and “Seg”. We model the attentional effect as a multiplicative gain according to a recent study using the same model (Rankin & Rinzel, 2022).

We fitted the tri-stability model to the percepts’ durations under varying conditions for individual subjects, contrasting with the typical approach of normalizing and pooling all data into a single idealized pseudo-subject. We considered several reasons. First, the pronounced individual differences observed in our tri-stability behavioral results undermine the reliability of a pooled data approach. Second, individualized model fitting provides further tests of the model’s explanatory power. Lastly, a quantitative description of individual variability could offer insights into the origins of these individual differences. To keep the fitting parsimonious, we adjusted only one parameter at a time, either noise or adaptation at a time. The outcomes suggest that both parameters are viable neural correlates for individual differences in multi-stable perceptions.

### 3. Model simulations reproducing the individuals’ and group’s key statistical characteristics

We fitted the noise (FN) or the adaptation (FA) parameter to align with each subject’s mean percept durations at varying Δ*f*s. The simulations with the optimized parameters reproduced the behavioral patterns decently. We evaluated the alignment between experimental data and model simulations using three interconnected but distinct metrics: the proportion of time, mean duration, and probability of occurring (Figure 4 and 5).

**Figure 4.**
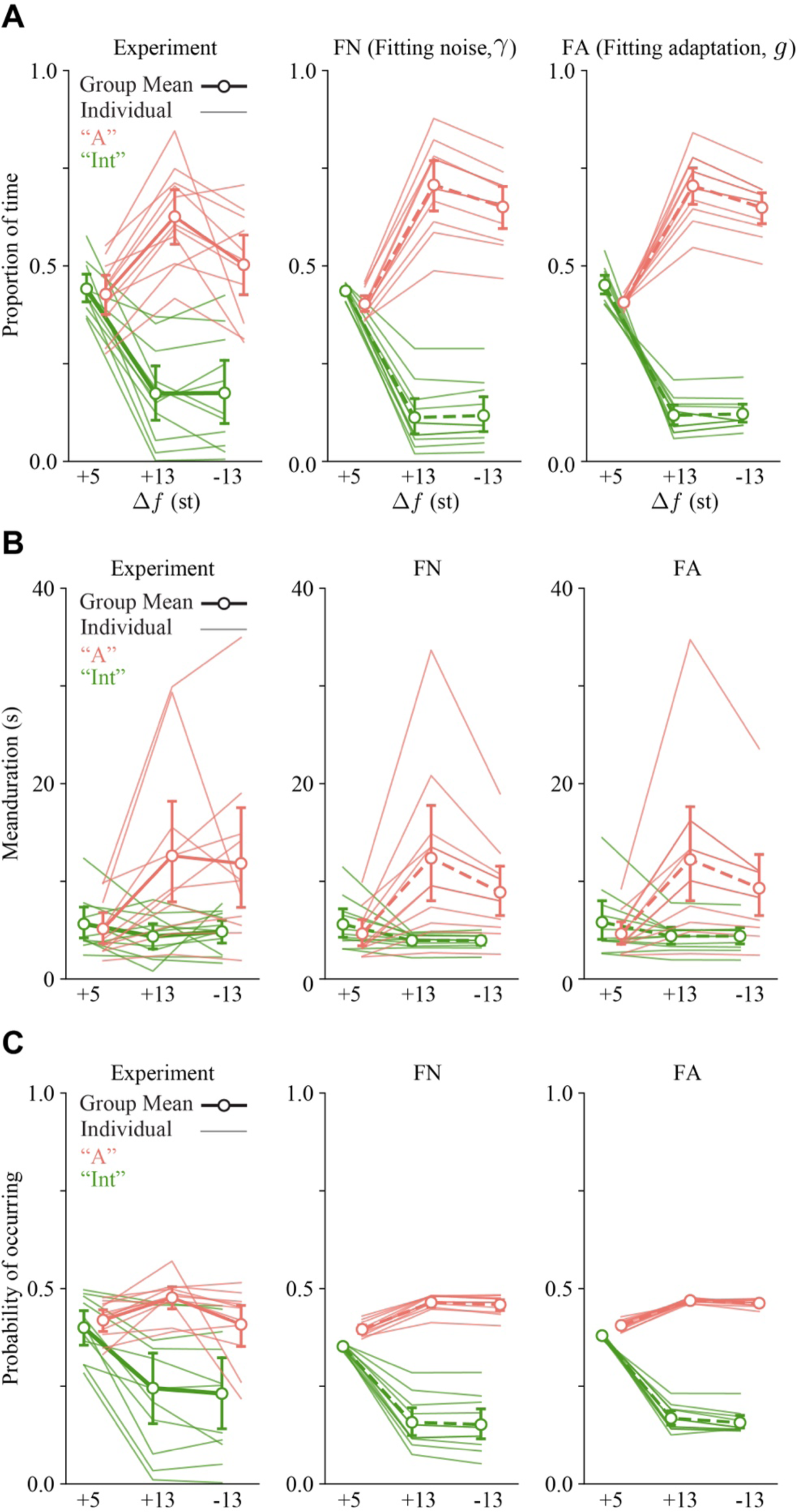
Model simulations reproduce the experimental results at an individual level. The left column shows the experiment results, the middle column shows the simulations using the model which assumes subjects differ in noise level (γ), and the right column shows simulations that assume subjects have different adaptation strengths (*g*). Fits with either parameter reproduce the average statistics (thick lines) equally well with differences regarding individual subjects (thin lines). Individual subjects’ fitted values of γ and *g* are shown in Figure 7. The same values of gamma and g were used in each panel for the simulations. The average values are computed from individual-based simulations. **A**, Proportion of time of “Int” percept and “A”. Participant-dependent fits on γ (middle column) show larger inter-subject variability than dependence on g (right column) despite their similarity in the group average. Notably, there are crossovers of individual lines in dependence on g between Δ*f* = +5 and Δ*f* = +13 for both “Int” and “A”, which are confirmed by statistical tests. However, such crossovers are not found in the experiment or in simulations dependent on noise. **B**, Mean duration of “Int” and “A”. The large individual differences are well captured by fitting with noise or with adaptation. **C**, Probability of percepts occurring. Fits with γ partially capture the individual differences while fits with g exhibit little variability. B percept is not plotted for clarity. Error bars represent 95% confidence intervals calculated from permutation.

**Figure 5.**
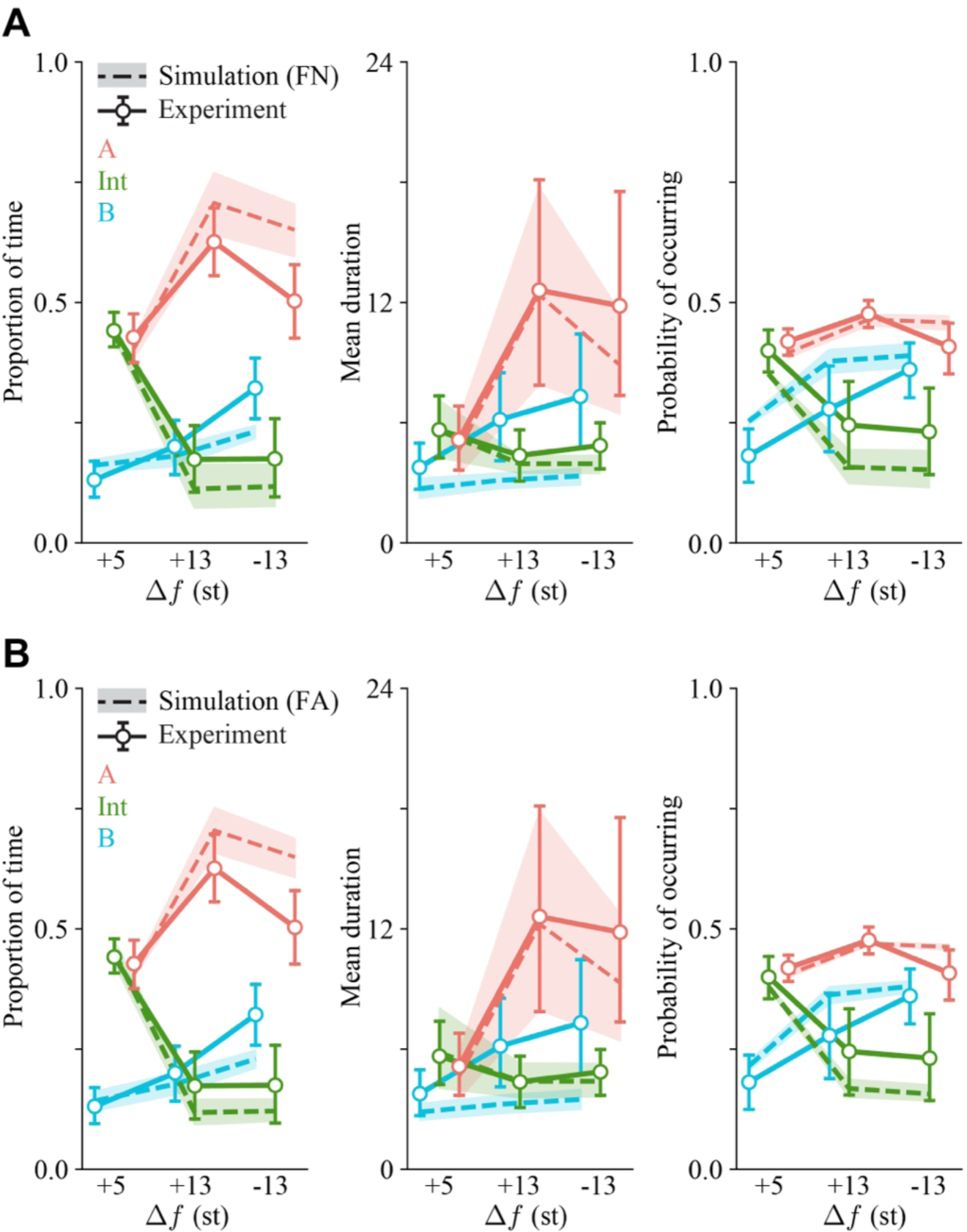
Direct Comparison of group-level statistics between the experiment and models. Both ways of parameter fitting (using γ or g) allow the model (dashed lines) to capture adequately the group-average summary statistics (proportion of time, mean duration, and probability of occurring) and the degrees of variability observed in the experiment (solid lines). Error bars (experiment) and shadings (simulation) represent 95% confidence intervals calculated from permutation.

The model simulations successfully replicated several key characteristics. In terms of the proportion of time (Figure 4A), the simulations pinpointed Δ*f* = +5 as the equidominance point between “Int” and “A”, identified a marked increase in the proportion of “A” and a decrease in “Int” when Δ*f* increased from +5 to +13, and demonstrated the effect of swapping tones’ frequencies, which slightly reduced “A” while leaving “Int” unchanged. Concerning the mean duration (Figure 4B), “Int” was relatively stable across subjects, whereas “A” was significantly extended at Δ*f* = ±13 versus Δ*f* = +5. The model also exhibited the trend that a percept with a longer average duration showed greater variability among individuals. Finally, trends in the probability of occurring paralleled those in the proportion of time but were less pronounced (Figure 4C). Notably, “A” showed more constrained variability than “Int”, yet “A” was more frequent, especially at higher Δ*f*s–another nuance captured by the simulations.

Both FN (Figure 4 middle column and Figure 5A) and FA (Figure 4 right column and Figure 5 B) reflected these dynamics, with FN capturing a wider range of intersubject differences than FA. Overall, both fitting strategies captured the general patterns and individual divergences across the metrics, albeit with some discrepancies: “B” percepts’ mean durations at Δ*f* = +13 were underestimated, and its probability of occurring at Δ*f* = −13 was slightly overpredicted. Although we fitted for different absolute values of Δ*f*s (5 and 13) separately, their best-fit values showed a strong positive correlation across subjects (Figure 6). This indicates the fitting can be further simplified by introducing a fixed scaling factor for different absolute values of Δ*f*. The range of best-fit γ is 0.057 ∼ 0.130 for Δ*f* = +5 and 0.043 ∼ 0.128 for Δ*f* = ±13. The range of best-fit *g* is 0.061 ∼ 0.215 for Δ*f* = +5 and 0.023 ∼ 0.216 for Δ*f* = ±13.

**Figure 6.**
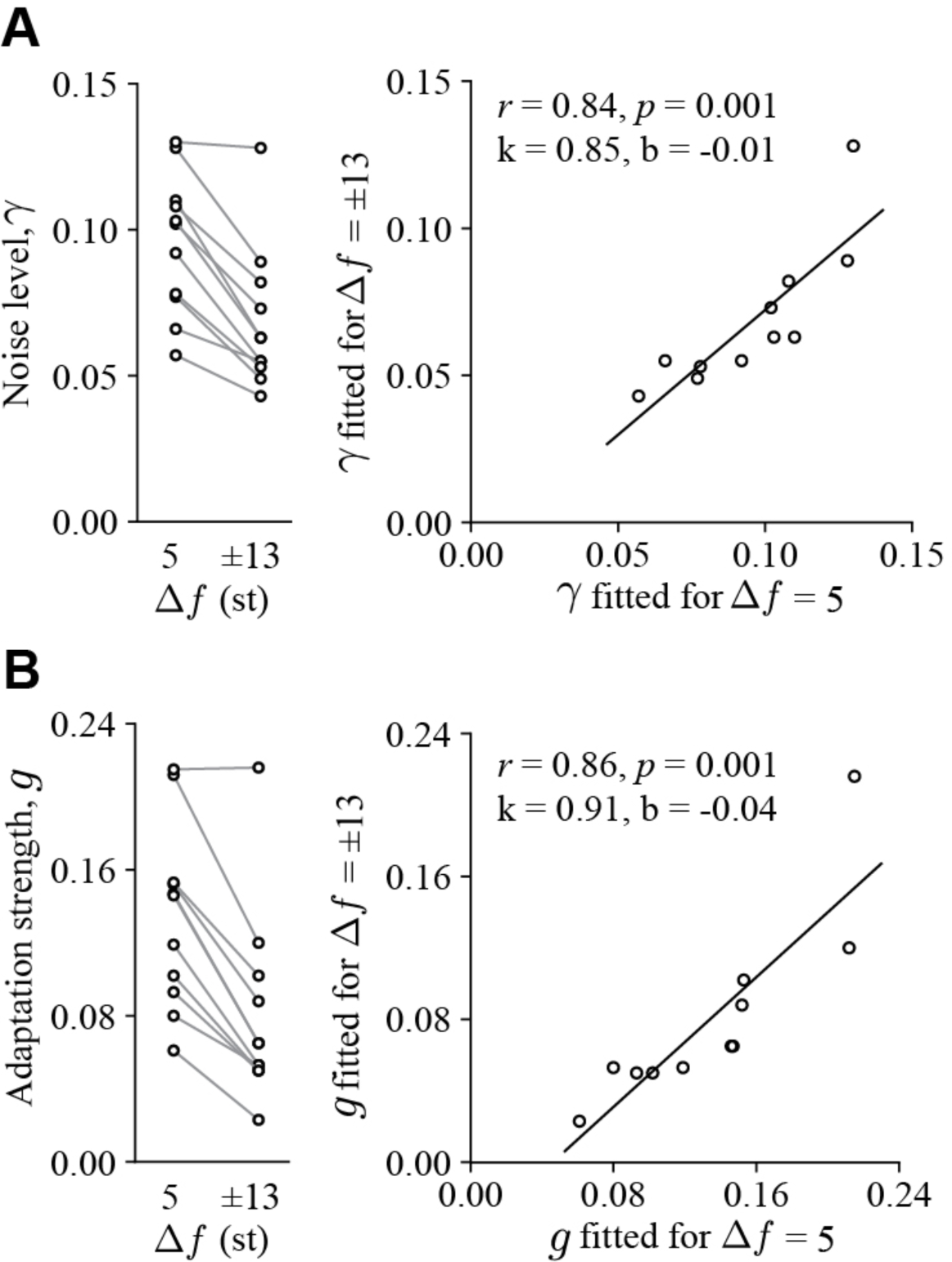
Individuals’ parameters fitted for different conditions follow a linear relation. Parameter γ or *g* is assumed to be dependent on the absolute value of Δ*f* and subjects, so parameters are fitted at an individual level respectively for Δ*f* = ±13 and Δ*f* = +5. Parameters γ and *g* are both smaller at Δ*f* = ±13 than at Δ*f* = +5 (left column) because the empirical mean durations are larger at Δ*f* = ±13 than at Δ*f* = +5. Post hoc analysis shows that parameters fitted for different absolute values of Δ*f* are highly correlated. **A**, Left: the noise levels parameter γ fitted for all eight subjects. Right: γs fitted for Δ*f* = ±13 are correlated with γs fitted for Δ*f* = +5 (*r* = 0.83, *p* = 0.010). Each pair of circles connected by a line represents a subject. **B**, Left: the adaptation strength parameter *g* fitted for all eight subjects. Right: *g*s fitted for Δ*f* = ±13 are correlated with γs fitted for Δ*f* = +5 (*r* = 0.90, *p* = 0.002). Each dot represents a subject.

In the final analysis, we compared our individual-based fitting approach with a group-mean-based fitting strategy. We juxtaposed the experimental results, the average results from individual simulations, and simulated results from the group-mean-fitted parameters (Figure 7). While the mean results from the individual models fell within the 95% confidence interval of the experimental results in most conditions, the group model often did not. Quantitatively, the average mean squared errors (MSEs) for the three indicators were generally lower in the individual models than the group-mean model (Figure 7, bar plots), except for the mean duration. Considering the model fitting error was computed solely over the mean duration, the other two indicators provide a clearer picture of the models’ true performance. In summary, fitting the model on an individual basis outperformed using the group averages. These findings provide a basis for future tests on noise and adaptation variation being neural correlates of individual differences in auditory streaming.

**Figure 7.**
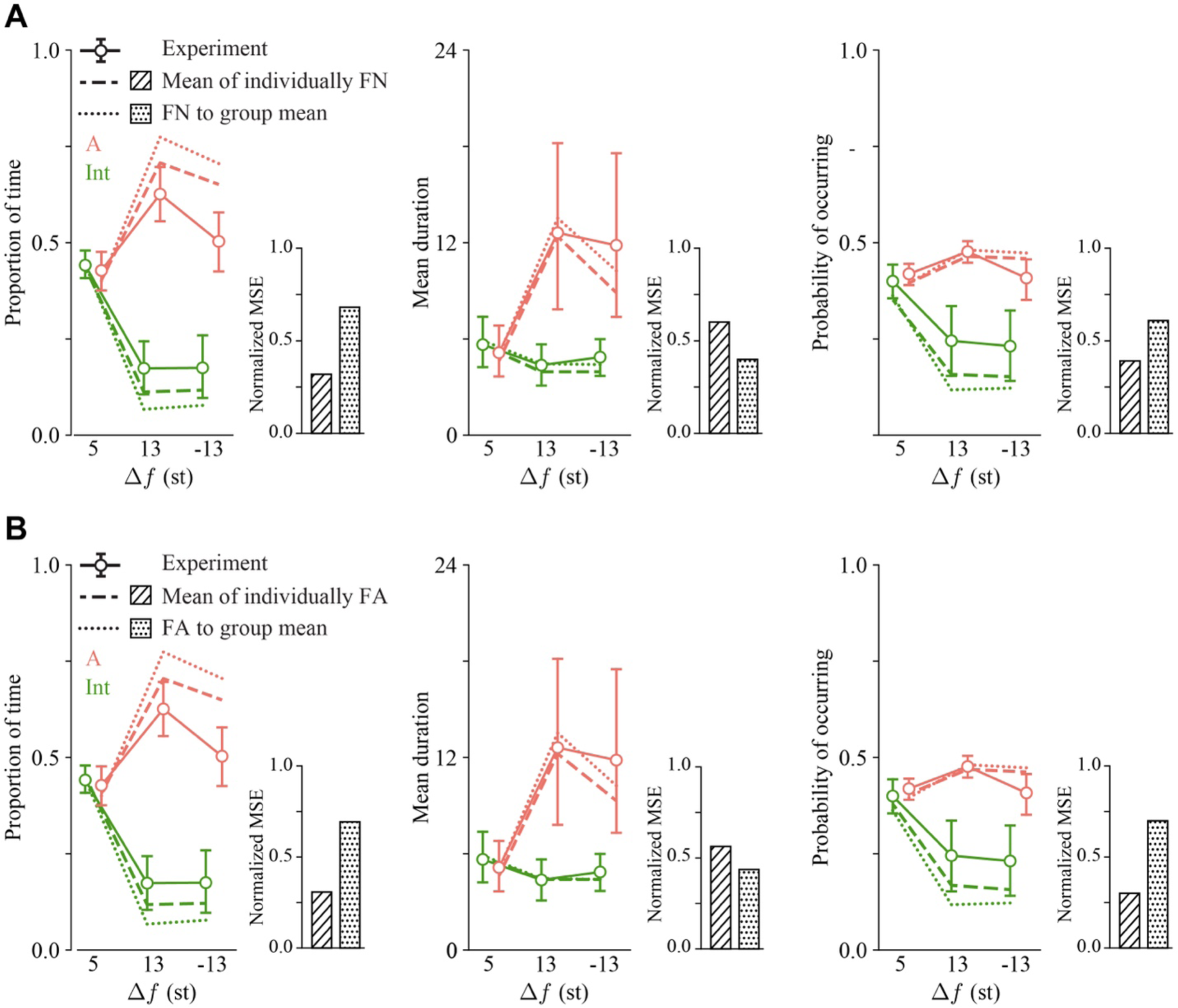
Fitting noise or adaptation to each individual outperforms fitting either parameter to the group mean. Comparison of the mean of individual models (dashed lines) and one model parameter fitted to the grand mean duration averaged across subjects (dotted line) with the experiment (solid line) – fits with noise level, γ (FN, **A**), and fits with adaptation strength, g (FA, **B**). The normalized mean squared errors of the two different parameter fits in the model are displayed to the right of each plot. Both the individual models and the group-mean model capture the fitting index well (mean duration, middle column), however, only the mean of the individual models falls into the 95% CI of the experimental results in the proportion of time (left column) and probability of occurring (right column). The normalized MSE of the single model (dotted patch) considerably exceeds the MSE of the mean of the individual fits in both the left (proportion of time) and right (probability of occurring) panel.

## Discussion

We reevaluated a neuro-mechanistic sensory model of multi-stable auditory perception by expanding its application from bi-stability to tri-stability. First, we collected data reflecting individuals’ perceptions that randomly switched among three interpretations during auditory streaming. The stimuli were triplets with an ABA_ABA_… pattern which admitted an integrated percept and two types of segregated percept–A or B sequence being in the foreground. We considered large Δ*f* to promote tri-stability and further test the model’s accountability. The considerable individual differences observed during the experiment posed a substantial challenge to reproduce the results with a single model. Nevertheless, we adopted this model without altering its fundamentals, given that has three competing units and inherent potential for tri-stability. The model proved a good fit for individual subjects’ mean percept durations, either by adjusting the noise parameter or the adaptation parameter. Additionally, the proportions of the three types of percepts and their frequencies of occurrence under varying conditions, though not included in model fitting, were reproduced by the simulation for free. Our results underscore the model’s adaptability and posit noise and adaptation as explanatory factors of the inter-individual variability observed in auditory tri-stability.

### Bi-modal subjects

Upon analyzing the histograms of percept durations for individual subjects, we found that four subjects were better fit with a bimodal distribution rather than a unimodal distribution for percepts at Δ*f* = +5 or ±13 (Supplementary Figure 3). Due to the insufficient commonality among the bimodal cases, we could not characterize the two modes beyond their existence. How bimodality may relate to task difficulty or tri-stability is an open question. When considering only the seven subjects characterized by unimodal distributions, the population histograms conformed to log-normal distributions (Supplementary Figure 5). However, when aggregated across all eleven subjects, the population histograms did not consistently adhere to a log-normal pattern (Supplementary Figure 4). This discrepancy between the seven and eleven subjects has been summarized in the condition-aggregated histograms depicted in Figure 3C. During model fitting, we treated the unimodal and bimodal subjects equally as the fitting was exclusively based on mean durations.

### Equidominance point in tri-stability

The distinction between perceptual tri-stability and bi-stability extends beyond merely adding a response option. First, the equidominance point shifts towards Δ*f* values smaller than the equidominance point (+5) reported in studies that offered only two response options (Rankin et al., 2015; Rankin & Rinzel, 2022). A previous study that incorporated comparable stimuli also showed that the equidominance point between segregation and integration would be smaller than Δ*f*=+5 in a tri-stability task (Hupé & Pressnitzer, 2012). However, such a trend in their results was less pronounced than ours, potentially because they used shorter presentation time (only 30 seconds), slightly smaller presentation rate (8.3 Hz), and the amalgamation of results from Δ*f* = ±3, ±5. Finally, we observed in our data a new type of equidominance between “Int” and “A” at Δ*f* = +5. Whether it’s a reliable pattern in auditory tri-stability or is specific to our setting and task requires future consideration.

### Noise and adaptation as general factors, not modality-specific, explaining individual differences in perceptual multi-stability

The large variability between subjects in auditory tri-stability, also observed by (Hupé & Pressnitzer, 2012), poses a significant challenge for model fitting. Despite this complexity, we fitted the model to replicate mean durations of distinct percepts, conditions, and subjects by only adjusting the noise or adaptation parameters. The simulations conducted with these fitted parameters turned out to reproduce acceptably the proportions of percepts and frequency of occurrences, too. Noise and adaptation are longstanding proposed driving influences in multi-stable perception (Winkler et al., 2012; Huguet et al., 2014; Meso et al., 2016). Our minimalist approach posits that these two factors could be essential in explaining the individual differences observed in auditory tri-stability.

Previous research has hinted at the existence of a partially modality-independent mechanism in perceptual multi-stability (Denham et al., 2018; Einhäuser et al., 2020), though no specific brain areas have been identified to date. Apart from that, strong inter-modality correlations across-subjects have been found between the visual and auditory domains. In this context, we put forth that noise and adaptation function as general individual traits, forming the foundation for the variations observed in perceptual multi-stability. This assertion gains additional support from a model-based study on tri-stable visual motion perception (Meso et al., 2016), where the results for individual subjects were effectively replicated by assuming their differences in the noise and a decision-making parameter.

### The role of attention in auditory multi-stability

We introduced an attentional gain parameter, denoted by ρ, to scale the total input directed to the “Int” unit in the competition stage. This was motivated by the observation that differentiating between the two segregated percepts, “A” and “B”, in tri-stability was more nuanced and challenging than distinguishing between integrated and segregated percepts in bi-stability. As a result, we suppose that the tri-stability task led participants to direct more attentional resources to “A” and “B” than to the “Int” percept. The notion that attention boosts the multiplicative gain of the inputs at the competition stage is supported by a study conducted by (Rankin & Rinzel, 2022). They investigated deliberate attentional control’s influence on auditory bi-stability. Within the same model framework, their behavioral results were most accurately described by assuming a static amplification of the inputs (from A1 responses) to the attended units/percepts at the competition stage.

Our incorporation of the task-induced attentional mechanism finds further support in a study comparing auditory tri-stability with visual tri-stability (Hupé & Pressnitzer, 2012). The researchers biased the perception towards one of the segregated percepts by adjusting the intensity of tone triplets in the auditory streaming or by introducing occlusion cues for visual plaids. They found that the inter-subject variability was more prominent in audition. To account for this difference, the researchers postulated that the dominance of either of the segregated auditory percepts involves attentional shifts. Meanwhile, the foreground-background separation in visual plaids is inherent and occurs naturally. This was evidenced in their experiments where significant amplification of either sound stream (A or B) did not completely abolish the dominance of the other stream. In contrast, visual cues, such as opaque left-moving plaids covering right-moving plaids rendered responses bi-stable, even though three options were provided.

### Tri-stability hypothesis regarding the build-up time course

Hupé and Pressnitzer (2012) linked tri-stability to the build-up phase. Their “tristability hypothesis” posits that the inertia of the initial “Int” percept, meaning its prolonged duration compared to subsequent “Int” percepts within a single presentation, is due to perceptual tri-stability. By incorporating occlusion cues, they successfully eliminated the first-percept inertia/prolongation in the perception of multi-stable visual motion. Moreover, in the ABA_ auditory streaming paradigm, they found a (marginally significant) negative correlation between the initial percept’s inertia and the degree that the segregated percept is biased to only one of the streams.

Our experimental findings are partially consistent with the tri-stability hypothesis. We observed first “Int” percept prolongation at Δ*f* = +5, but not at Δ*f* = ±13, as depicted in Supplementary Figure 7. This held true even though tri-stability was largely preserved at larger Δ*f*s. The breakdown of first-percept prolongation as Δ*f* increases was also observed in Nguyen et al. (2020) at Δ*f* = 7. Furthermore, “first percept bias” is a distinctive concept compared with “first percept inertial”. The former suggests that the initial percept is more likely to be integrated than segregated. All tested conditions (Δ*f* = +5, ±13) exhibited a build-up function that gradually increased from near-zero and plateaued at around eight seconds (Supplementary Figure 8). The pattern echoes previous findings and affirms the reliability of our data.

### The order of the tone frequencies

Whether transitioning from low to high to low, or vice versa–was often taken as of no consequence in studies on bi-stability. However, two studies concerning tri-stability (Denham et al., 2014; Thomassen et al., 2022) uncovered distinct behavioral patterns when contrasting high-low-high and low-high-low triplets. Denham et al. (2014) suggested that specific tone frequencies influence the dominance pattern via perceptual saliency. To verify this conjecture, we contrast conditions of Δ*f*=+13 (high-low-high) with Δ*f*=-13 (low-high-low). Switching the frequencies of the A and B tones exhibited a similar but less dramatic effect in our results compared to (Denham et al., 2014). Specifically, the proportion of “A” decreased slightly, and the proportion of “B” only increased mildly from Δ*f*=+13 to Δ*f*=-13 in our experiment.

In our model, the tones’ perceptual saliency is reflected by their response amplitudes in A1. For simplicity, we introduced the relative saliency parameter, denoted as *q*, to denote the amplitude ratio of the B tone to the A tone. Rather than assuming an arbitrary connection between perceptual saliency and tone frequency, we constrained their relationship using the equal-loudness contours as a reference. Given that the curves depicting equally loud tones within the intensity-frequency space are not monotonic, we placed strategically the frequencies of the tones for different Δ*f*s (Supplementary Figure 6), to ensure the high-pitch tone was consistently louder (and therefore more salient) than the low-pitch tone in any pair. In other words, *q* < 1. The specific value of the parameter *q* was optimized at the group level. The use of equal-loudness contours is justified by the understanding that A1 responses reflect loudness (subjective intensity) in a linear fashion instead of objective intensity (Röhl & Uppenkamp, 2012)

### Models of perceptual multi-stability

Our model aligns with dynamical systems approaches, embodying biased competition, as developed by Laing & Chow (2002) and Wilson (2003). This approach has been extended by the works of Shpiro et al. (2007, 2009), Moreno-Bote et al. (2007), Huguet et al. (2014), and Meso et al. (2016), with a focus on elucidating the contributions of noise and adaptation in accounting for statistics. Notably, Moreno-Bote et al. (2007) and Huguet et al. (2014) distinguished noise-driven vs. adaptation-driven switching.

Along this line, our approach extends neuro-mechanistic modeling within the context of auditory bi-stability, as developed by Rankin et al. (2015). Apart from noise and adaptation, alternative driving factors have been proposed, including excitation and inhibition by Shpiro et al. (2007), and attention in binocular rivalry by Li et al. (2017). Considering the diversity of possible perceptions during auditory streaming, both Mill et al. (2013) and Ferrario & Rankin (2021) proposed models incorporating an initial stage of identifying and forming different perceptual interpretations, preceding competition. Furthermore, Little et al. (2020) put forth a hierarchical model with features of varying time scales and competition occurring across all levels.

A different category of models for bi-stability hinges on evidence accumulation, whereby perceptual switching is triggered once the representation of evidence against the current percept surpasses a threshold level (Barniv & Nelken, 2015; Cao et al., 2016; Nguyen et al., 2020). Certain models within this category, specifically those by Cao et al. (2016) and Nguyen et al. (2020), employ a discrete collection of binary units to support the opposing evidence. Cao et al. (2016) highlighted the scaling properties across various orders of statistics. Further, their recent paper (Cao et al., 2021) drew attention to the serial dependence in sequential percepts: the durations of successive percepts are positively correlated. Their model replicated this positive correlation. Here, we have sought primarily to determine whether the summary first-order statistics in tri-stability can be succinctly reproduced by our model (Rankin et al., 2015) within the noise-adaptation framework. Whether our model could further explain serial dependency remains a topic for future exploration.

Some earlier computational approaches to auditory scene analysis including stream formation, not necessarily for multi-stable alternations, were described and compared by Wang & Brown (2006a) (from Wang & Brown (2006b), that includes several other approaches). Among the models covered were those of (McCabe & Denham, 1995) (featuring spectrally-based multi-channel competitive interactions, influenced by previous activity, to represent the temporal development of streams); Wang & Brown (1999), Wang & Terman (1995) (a dynamical systems approach using oscillation correlations as a basis for auditory perceptual grouping); and Grossberg et al. (2004) (proposing a resonance between top-down expectation and bottom-up harmonic filters, that adaptively leads to perceptual grouping).

### In closing and looking ahead

We have shown that a competition framework for perceptual bi-stability accounts well also, as an emergent property, for the dynamics of tri-stability. Our model (Rankin et al., 2015) was not developed with tri-stability in mind but rather to provide a neuromechanistic description of perceptual alternations. In actuality, the experiments reported here were carried out to test our prediction based on the model simulations of tri-stability. Our formulation of the competition model as *sensory-based,* taking A1-responses (Fishman et al., 2001) to tone-sequence stimuli as inputs to three tonotopically organized units, was key in enabling the emergent property. In contrast, our earlier model for visual tri-stability (Huguet et al., 2014) was designed with *percept-based inputs* and competition between three *percept-specific units*. Looking ahead, we are motivated to ask if the current model, with some extension, may account for other perceptual patterns, say, like some of those offered as choices to subjects listening to triplets: AB-- and -BA-(Denham et al., 2014). In Ferrario & Rankin (2021) an intermediate stage (think of secondary auditory cortex, A2), between A1 and a downstream competition stage, was proposed in which pattern types for sequential inputs ABABAB… were identified by applying dynamical systems methods on competition amongst tone-selective units that were mutually coupled with excitation and delayed inhibition. The various attractor states were proposed to compete downstream, ultimately giving rise to alternation dynamics. This intermediate stage can be thought of as providing an alternative to dynamic object identification amongst a library of patterns as in Mill et al (2013) or the combined effects of pre-Object stages in Little et al (2020). The determination of patterns as elemental short tone groupings can bring together rhythmic pattern formation and auditory streaming as suggested in Ferrario & Rankin (2021). A further extension in the spirit of neuromechanistic modeling could expand the input representation, allowing for broader spectrally distributed sound combinations. Rather than two pure tone sequences a model with a continuously distributed frequency axis for A1 processing (e.g., Levy & Reyes, 2011) could open consideration of competition, for example, say, between sequences of harmonically-stacked tone patterns (Deike et al., 2012). In studying multi-stable visual perception, models with distributed sensory features have been developed and applied to describe, for example, binocular rivalry between gratings as represented on a ring network of orientation-selective neuronal subgroups (Laing & Chow, 2002) or spatiotemporal dynamics during binocular rivalry (Kang, 2009; Kang & Blake, 2010; Bressloff & Webber, 2012), and for tri-stable perception as for the barber pole motion stimulus (Meso et al., 2016).

## Supplemental Information

**Supplementary Figure 1.**
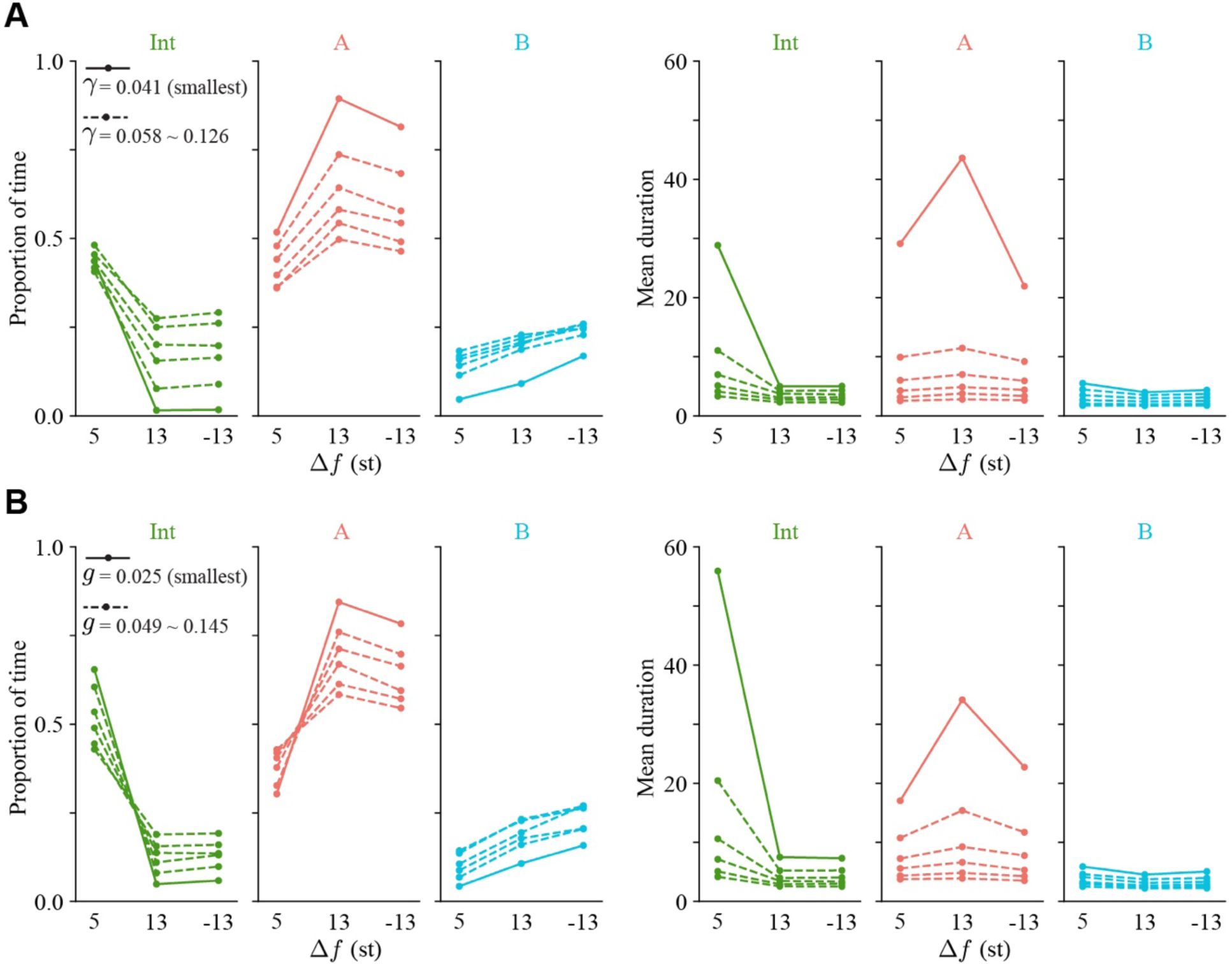
The model suggests that either the noise level or the adaptation strength can modulate the proportion of time and mean durations differently. **A**, the proportion of time (left) and the mean duration (right) of the three percepts as the noise level γ traversing a set of evenly spaced values (0.041 ∼ 0.126), with the solid line having the smallest γ. The proportion of time and mean duration of “A” decrease as gamma increases. For “Int” and “B”, the proportion of time increases with γ, but the mean duration decreases with gamma. Note that the mean duration of A (right in red) is especially sensitive to γ. B, the same indicators when the adaptation strength *g* traversing a set of evenly spaced values (0.025 ∼ 0.145). While the mean duration (right) shares a similar pattern as varying γ in A, the pattern of the proportion of time (left) is different. For “Int” and “A”, the proportions of time at Δ*f* = +5 move in the directions opposite to themselves at Δ*f* = ±13 (crossover of the lines).

**Supplementary Figure 2.**
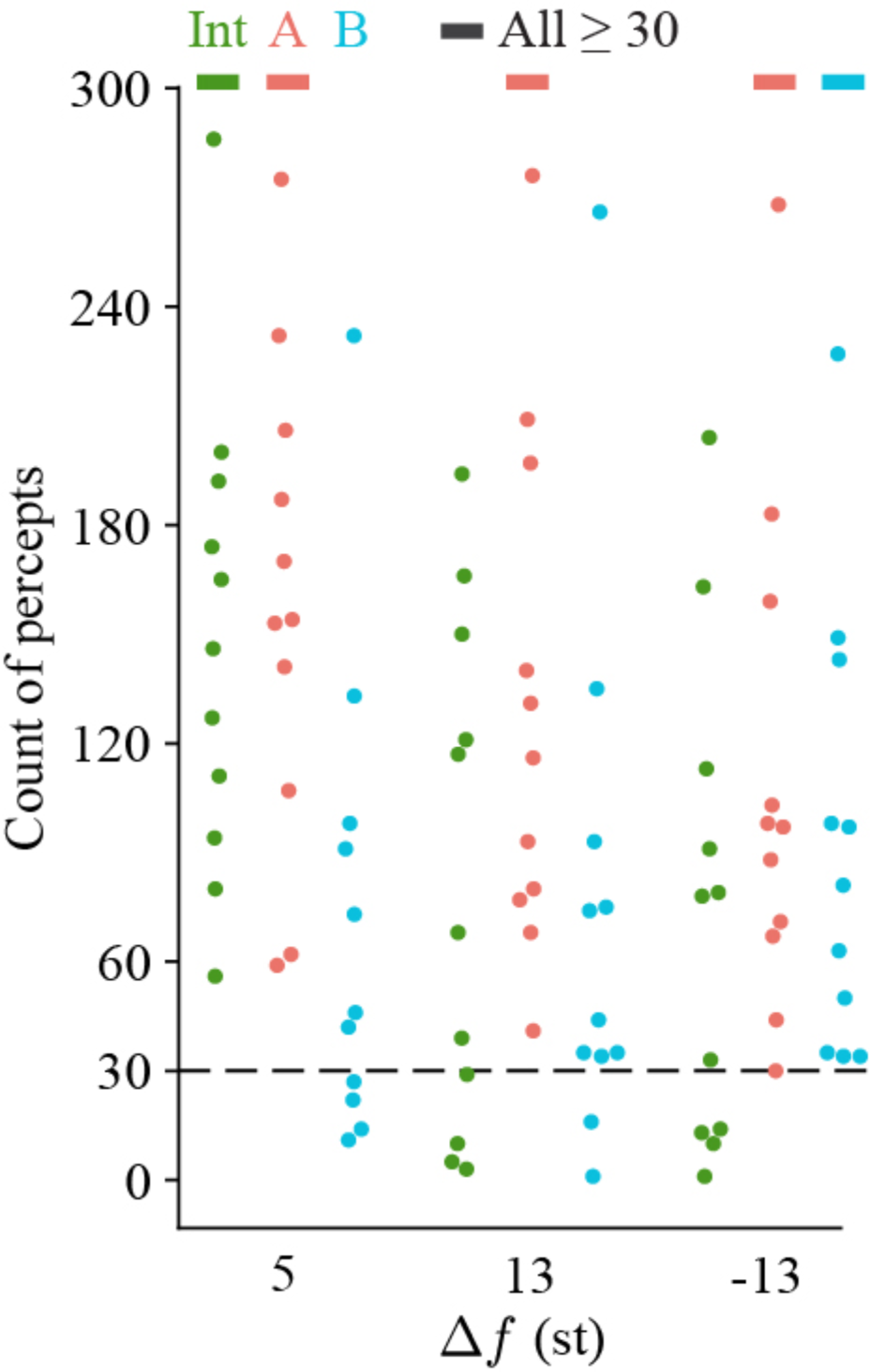
Conditions and percept types in which all subjects yield above 30 occurrences are used for model fitting. Each dot represents one subject under one condition. Solid shadings indicate the Δ*f*-percept conditions used in model fitting.

**Supplementary Figure 3.**
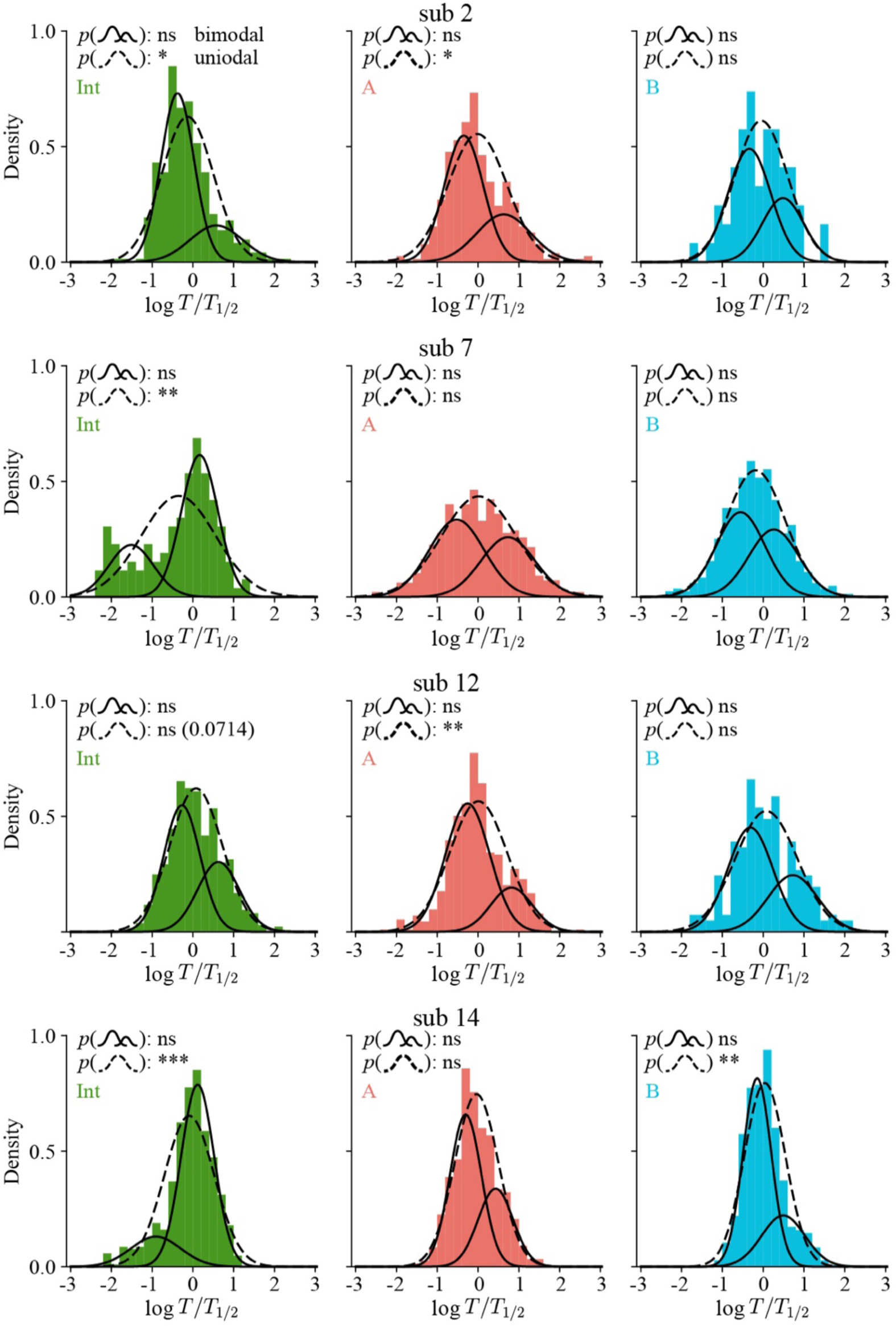
The duration histograms of four subjects for percept “Int”, “A”, and “B” in log scale, aggregated over Δ*f* conditions, are fitted by a bimodal Gaussian mixture model (solid) and a unimodal Gaussian (dashed). Each subject has at least one histogram that is better captured by the bimodal distributions than the unimodal distribution. We normalized the durations by their medians within each Δ*f* (+5, ±13), percept (“Int”, “A”, “B”), and subject (11 in total). The x-axes are of log scale. * indicates *p* < 0.05, ** *p* < 0.01, *** *p* < 0.001, and **** *p* < 0.0001.

**Supplementary Figure 4.**
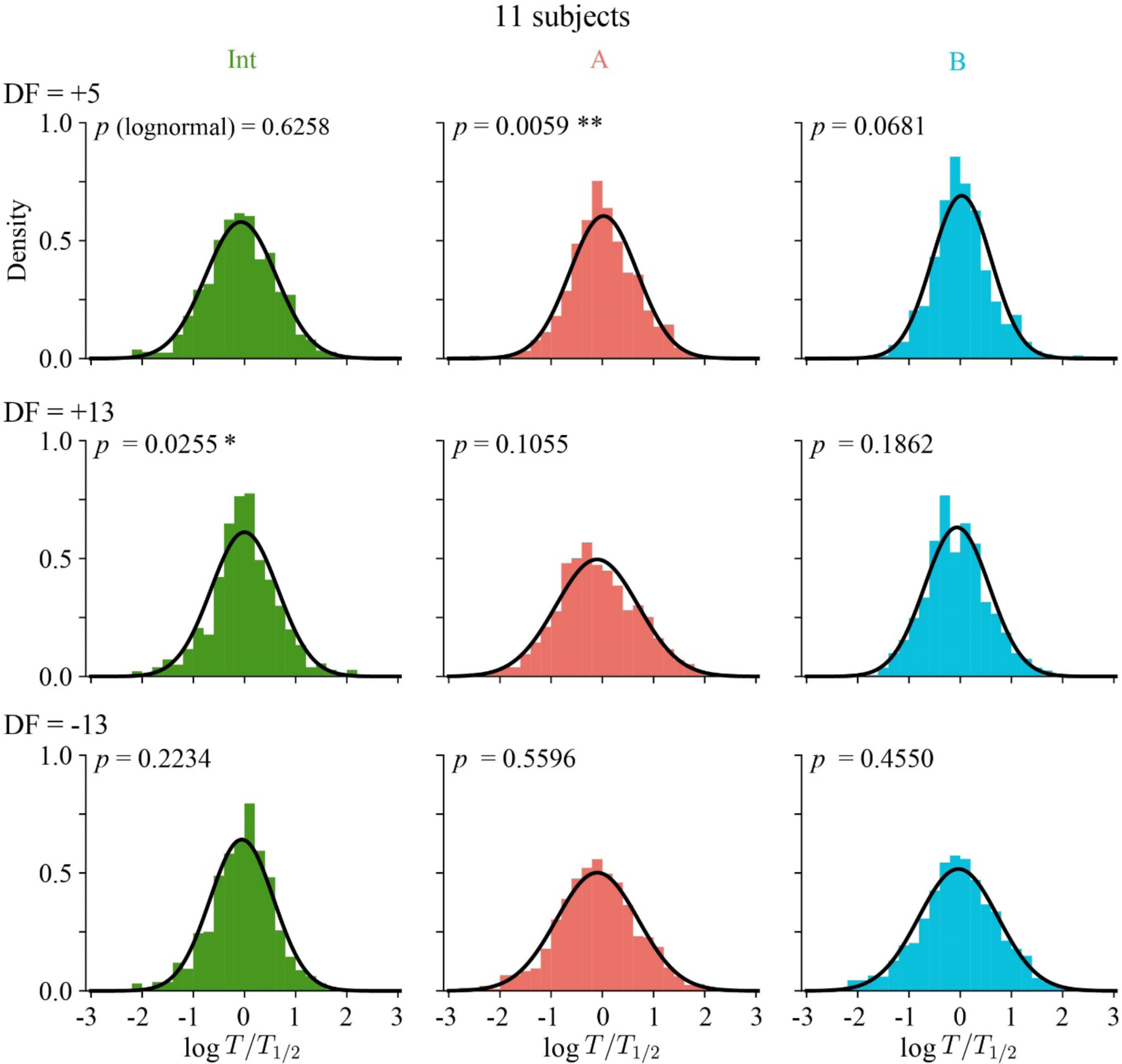
Histograms of percept dominance durations aggregated across all 11 subjects. Some of the histograms are not log-normally distributed because of the 4 bi-modal subjects shown in Supplementary Figure 1. The percept types (“Int” and “A”) exhibiting deviation from lognormality coincide with those in Supplementary Figure 1. The rows and columns correspond to different conditions and percept types respectively. * indicates *p* < 0.05, ** *p* < 0.01, *** *p* < 0.001, and **** *p* < 0.0001.

**Supplementary Figure 5.**
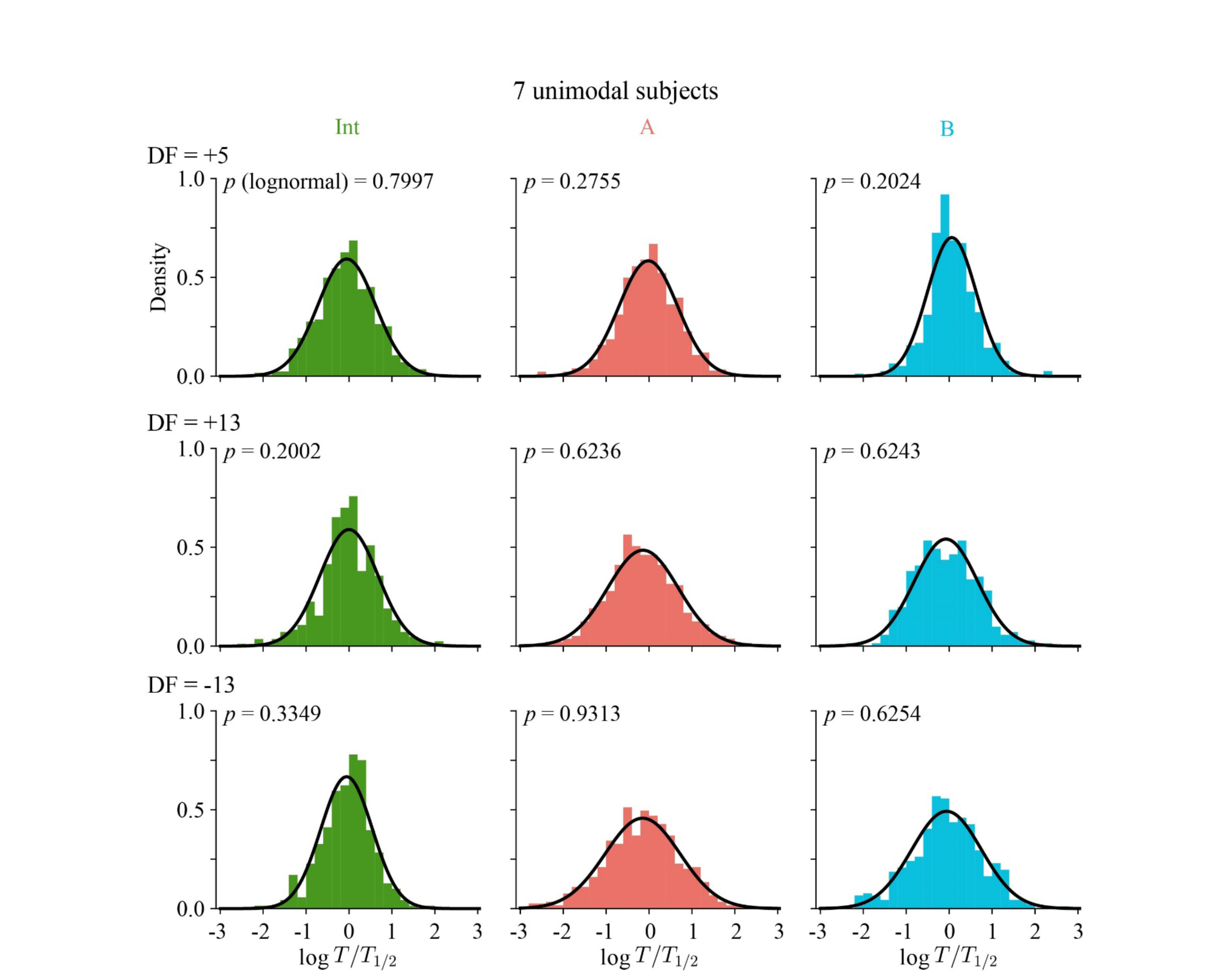
Histograms of percept dominance durations aggregated across the 7 unimodal subjects are all lognormal. Removing the four bimodal subjects shown in Supplementary Figure 1 reveals that the aggregated durations are lognormal distributed. The rows and columns correspond to different conditions and perceptual types respectively. * indicates *p* < 0.05, ** *p* < 0.01, *** *p* < 0.001, and **** *p* < 0.0001.

**Supplementary Figure 6.**
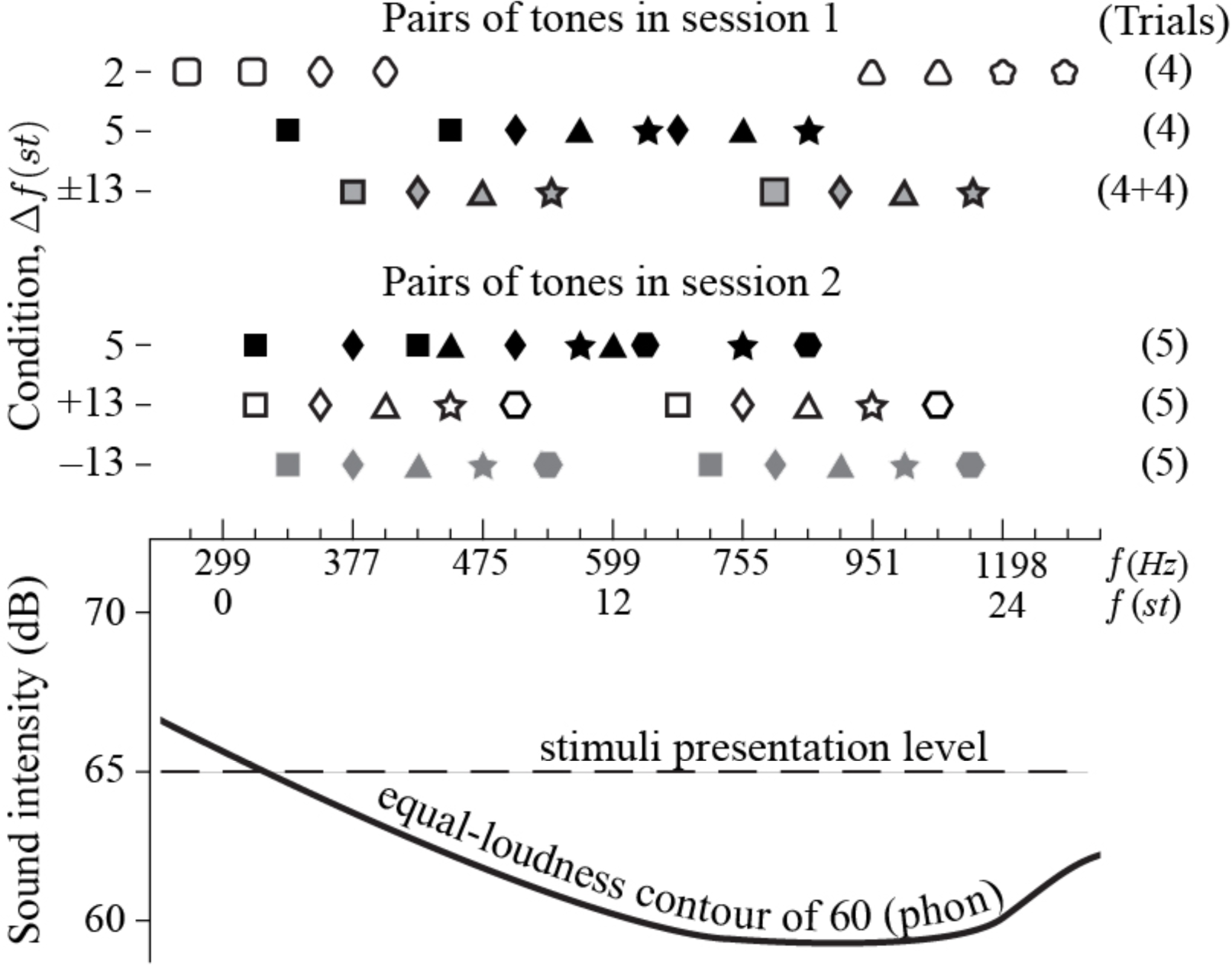
The pitches of the two tones used in all trials are selected to minimize repetition across trials and to constrain tone saliency (based on the equal-loudness contours). Upper half: arrangements of the tones’ pitches. Each like-symbolled pair (unique combination of shape, outline, and fill) in a row represents the two tones used in a trial, whose abscissas indicate the pitches in Hz and semitones (st). Session 1 (rows 1-3) has four conditions (Δ*f* = +2, +5, ±13) and Session 2 (rows 4-6) has three conditions (Δ*f* = +5, ±13). Δ*f* = +13 means that A tone is higher than B tone by 13 st while Δ*f* = −13 means that the A tone is lower in pitch than the B tone by 13 st. Lower half: the solid line is a section of the equal-loudness contour (ISO226) at 60 phon and the dashed line is the intensity (65 dB) at which all tones are presented. Tones along the equal-loudness contour are on average equally loud to the population. Hence, tones sitting on the left slope of the curve require larger intensity to be heard as loud as tones located at the basin. Such arrangements of stimuli ensure that the higher-pitch tone in a trial is on average louder and more salient than the lower-pitch tone. The dashed line shows how to link a tone to its pitch.

**Supplementary Figure 7.**
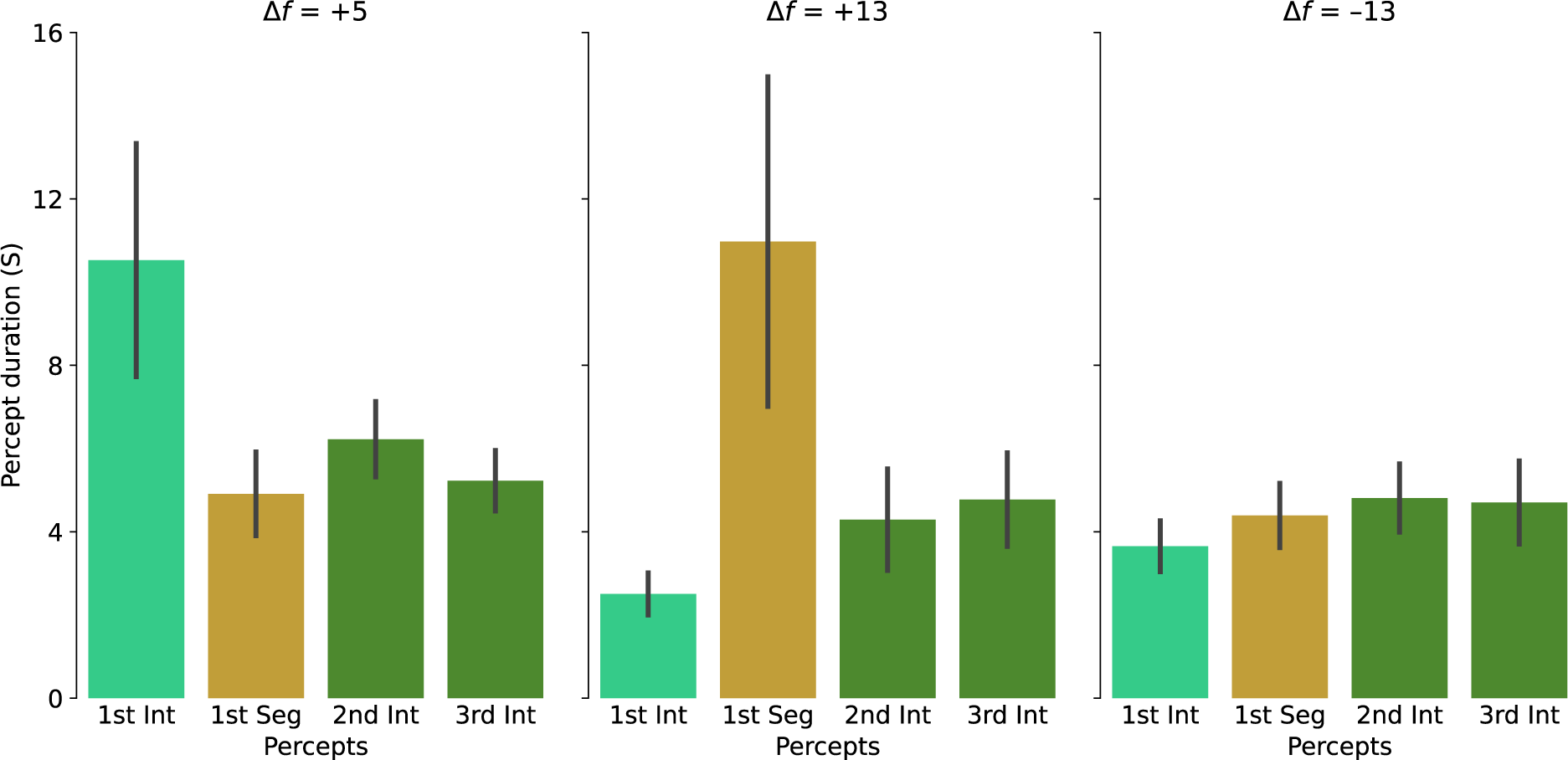
First percept inertia (prolonged “Int” duration) is only seen in Δ*f* = +5.

**Supplementary Figure 8.**
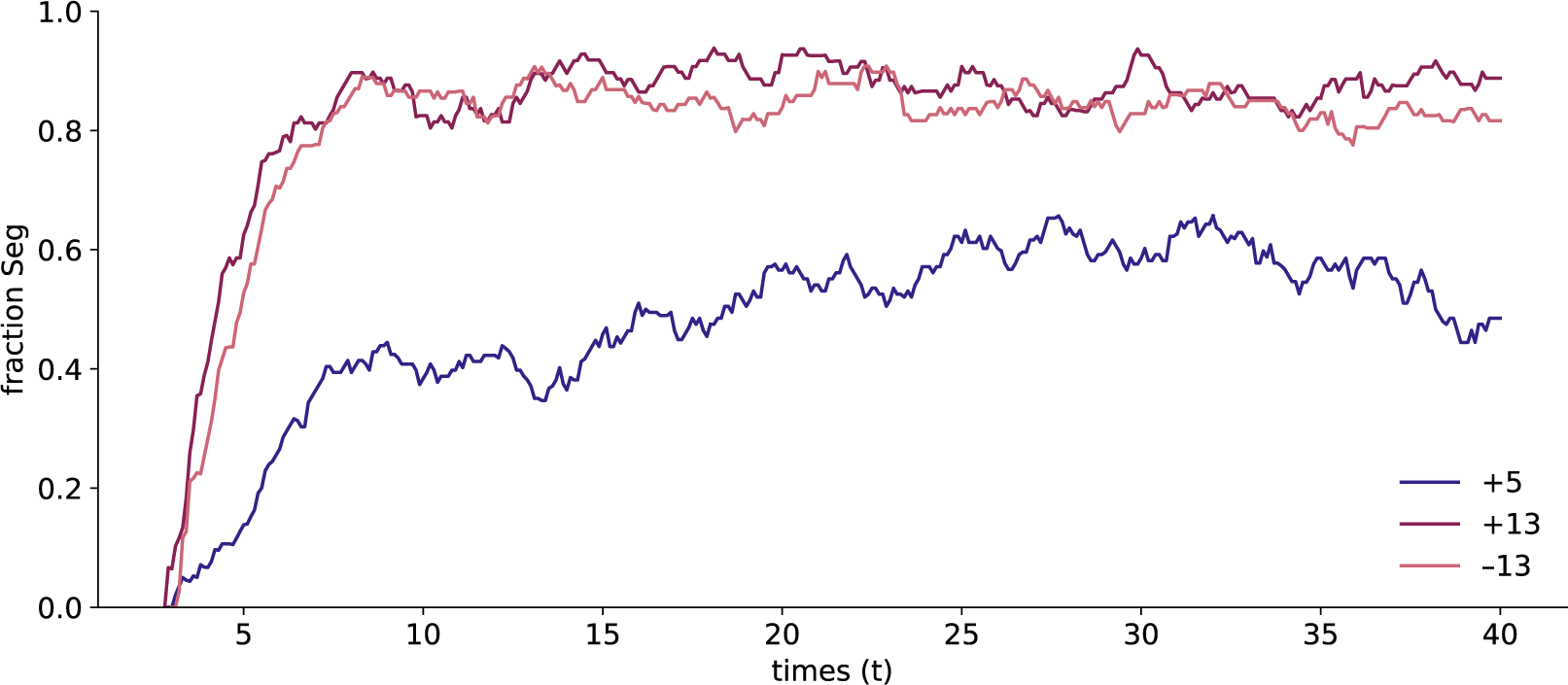
Build-up functions.

**Supplementary Table 1.**
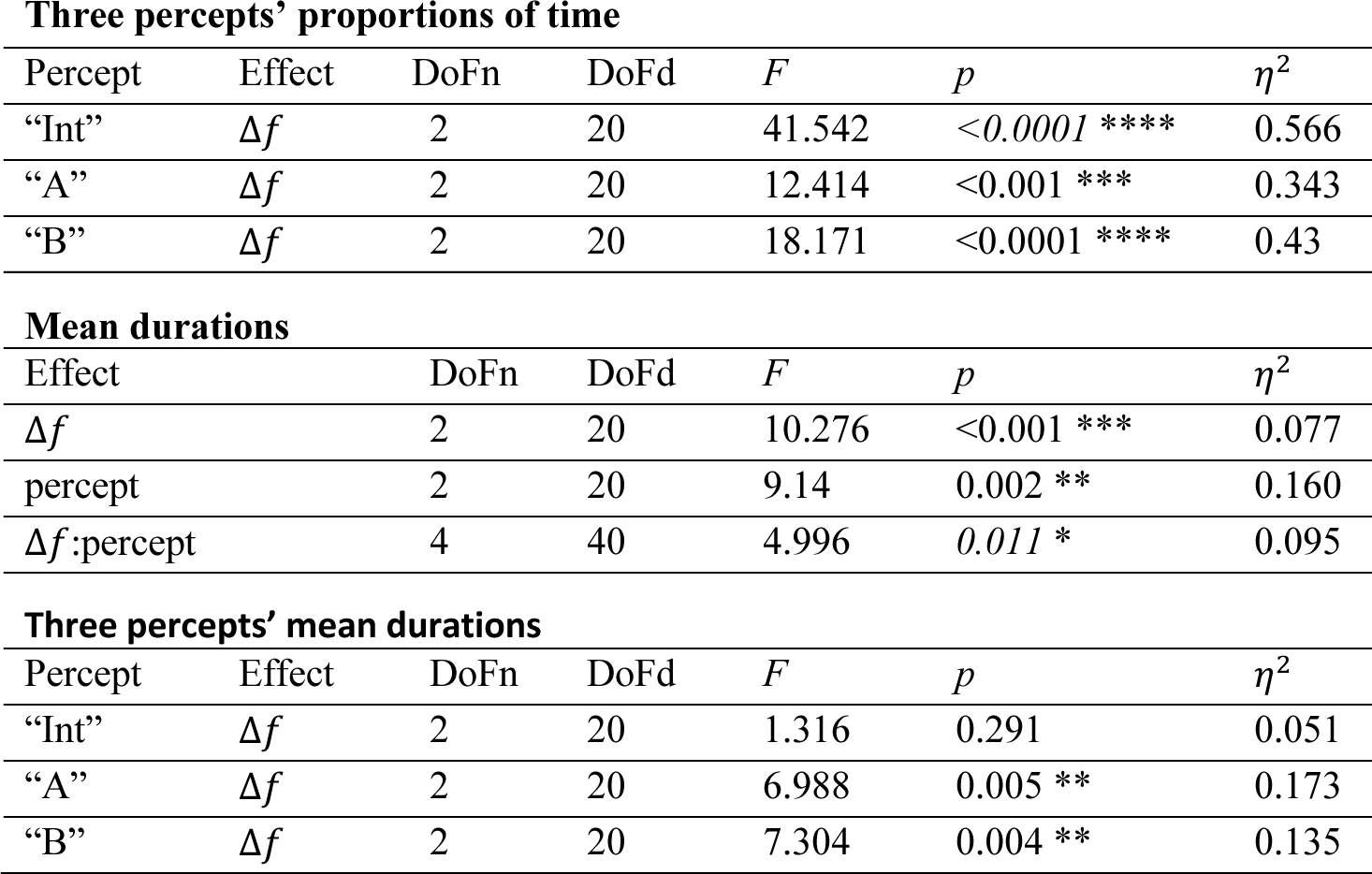
Repeated measures ANOVA table for proportions of time and mean durations. For each of the three percepts (“Int”, “A”, “B”), we applied a one-way ANOVA to the proportions of time with the tones’ difference in pitch Δ*f* (+5, ±13) as a within-subject factor. We applied a two-way ANOVA to the mean durations with Δ*f* and percept as within-subject factors. Additionally, for each of the three percepts we applied a one-way ANOVA to the mean durations with Δ*f* as a within-subject factor. We also showed the degrees of freedom (DoF), *F* static, *p* value, and generalized η^2^. We reported and italicized the Greenhouse-Geiser-corrected p values when the Mauchly’s test for non-sphericity reached significance. * indicates *p* < 0.05, ** *p* < 0.01, *** *p* < 0.001, and **** *p* < 0.0001.

Three percepts' proportions of time
| Percept | Effect | DoFn | DoFd | *F* | *p* | $\eta^2$ |
| --- | --- | --- | --- | --- | --- | --- |
| "Int" | $\Delta f$ | 2 | 20 | 41.542 | <0.0001 **** | 0.566 |
| "A" | $\Delta f$ | 2 | 20 | 12.414 | <0.001 *** | 0.343 |
| "B" | $\Delta f$ | 2 | 20 | 18.171 | <0.0001 **** | 0.43 |

Mean durations
| Effect | DoFn | DoFd | *F* | *p* | $\eta^2$ |
| --- | --- | --- | --- | --- | --- |
| $\Delta f$ | 2 | 20 | 10.276 | <0.001 *** | 0.077 |
| percept | 2 | 20 | 9.14 | 0.002 ** | 0.160 |
| $\Delta f$:percept | 4 | 40 | 4.996 | 0.011 * | 0.095 |

**Supplementary Table 1 Repeated measures ANOVA table for proportions of time and mean durations.**
| Percept | Effect | DoFn | DoFd | *F* | *p* | $\eta^2$ |
| --- | --- | --- | --- | --- | --- | --- |
| "Int" | $\Delta f$ | 2 | 20 | 1.316 | 0.291 | 0.051 |
| "A" | $\Delta f$ | 2 | 20 | 6.988 | 0.005 ** | 0.173 |
| "B" | $\Delta f$ | 2 | 20 | 7.304 | 0.004 ** | 0.135 |

**Supplementary Table 2.**
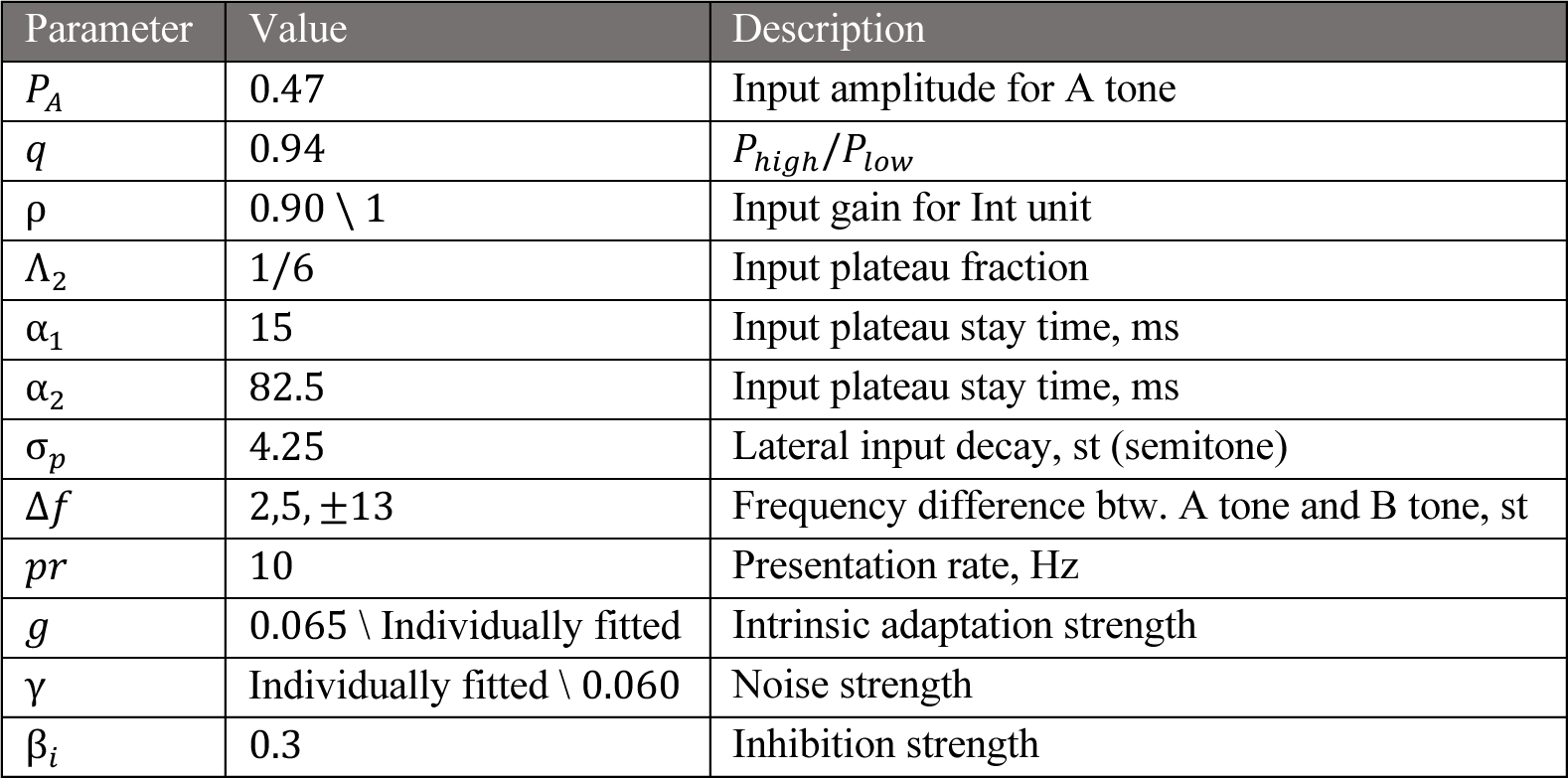
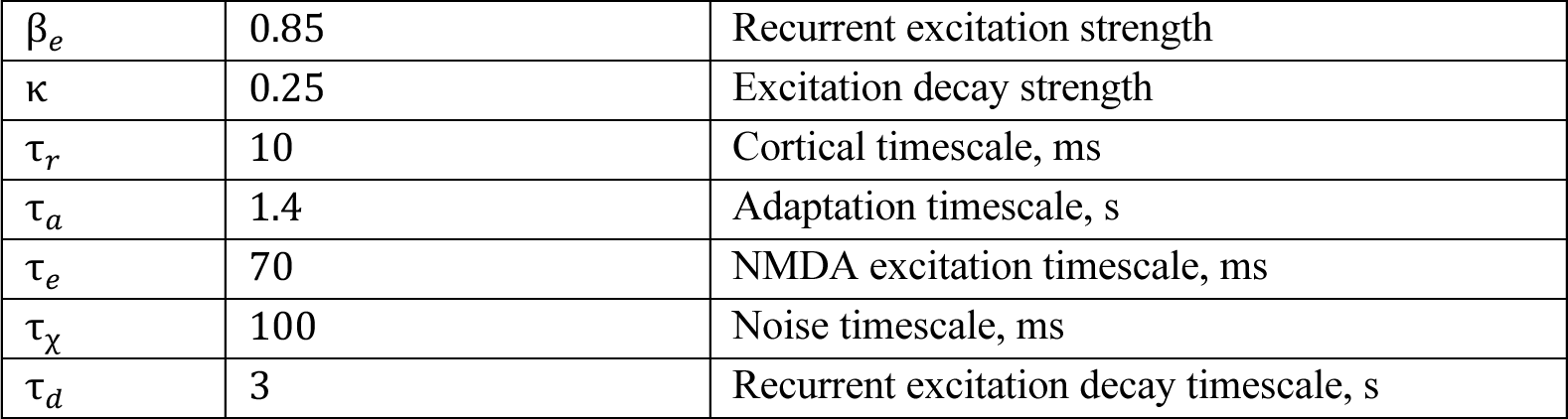
Model parameters. When we fitted model 1 for individual subjects, we set the noise level (γ) to be the free parameter while fixing the input gain of the Int unit (*q*_$%&_) at 0.9 and the adaptation strength (*g*) at 0.065. When fitting model 2 for individual subjects, we set *g* to be the free parameter while fixing *q*_$%&_ at 1.0 and γ at 0.06.

## Model Equations

We use the over-dot notation for the time derivative

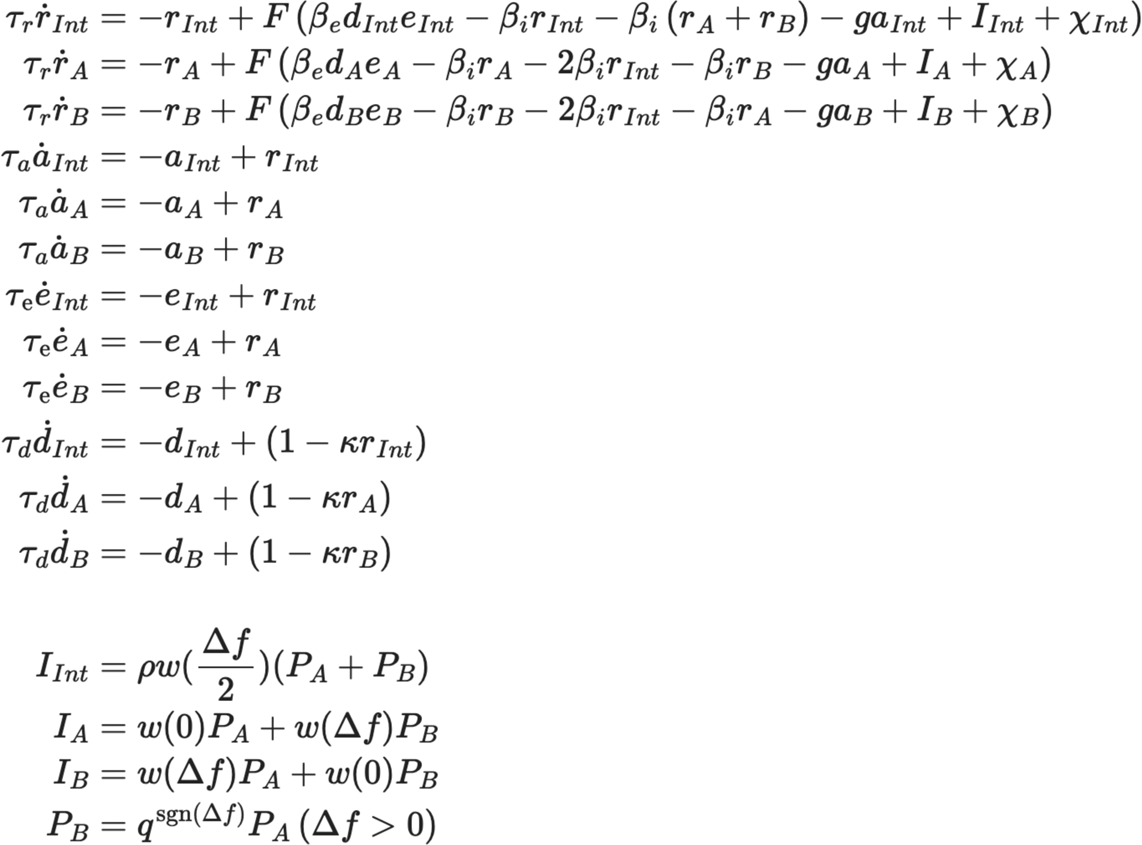

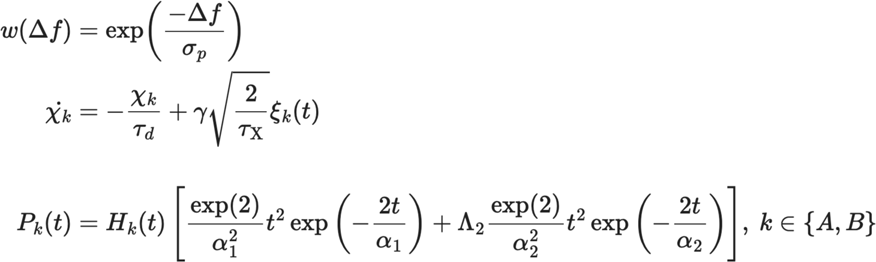

## Notes

### Competing Interest Statement

The authors have declared no competing interest.

